# An increase in reactive oxygen species underlies neonatal cerebellum repair

**DOI:** 10.1101/2024.10.14.618368

**Authors:** Anna Pakula, Salsabiel El Nagar, N. Sumru Bayin, Jens Bager Christensen, Daniel N. Stephen, Adam James Reid, Richard Koche, Alexandra L. Joyner

**Affiliations:** Developmental Biology Program, Sloan Kettering Institute, New York, NY, USA; Gurdon Institute, Cambridge University, Cambridge, UK; Department of Physiology, Development and Neuroscience, Cambridge University, Cambridge, UK; Center for Epigenetics Research, Memorial Sloan Kettering Cancer Center, New York, NY; Biochemistry, Cell and Molecular Biology Program and Neuroscience Program, Weill Cornell Graduate School of Medical Sciences, New York, NY, USA

**Keywords:** Nestin-expressing progenitors, NEPs, granule cell progenitors, ROS, regeneration

## Abstract

The neonatal mouse cerebellum shows remarkable regenerative potential upon injury at birth, wherein a subset of Nestin-expressing progenitors (NEPs) undergoes adaptive reprogramming to replenish granule cell progenitors that die. Here, we investigate how the microenvironment of the injured cerebellum changes upon injury and contributes to the regenerative potential of normally gliogenic*-*NEPs and their adaptive reprogramming. Single cell transcriptomic and bulk chromatin accessibility analyses of the NEPs from injured neonatal cerebella compared to controls show a temporary increase in cellular processes involved in responding to reactive oxygen species (ROS), a known damage-associated molecular pattern. Analysis of ROS levels in cerebellar tissue confirm a transient increased one day after injury at postanal day 1, overlapping with the peak cell death in the cerebellum. In a transgenic mouse line that ubiquitously overexpresses human mitochondrial catalase (mCAT), ROS is reduced 1 day after injury to the granule cell progenitors, and we demonstrate that several steps in the regenerative process of NEPs are curtailed leading to reduced cerebellar growth. We also provide preliminary evidence that microglia are involved in one step of adaptive reprogramming by regulating NEP replenishment of the granule cell precursors. Collectively, our results highlight that changes in the tissue microenvironment regulate multiple steps in adaptative reprogramming of NEPs upon death of cerebellar granule cell progenitors at birth, highlighting the instructive roles of microenvironmental signals during regeneration of the neonatal brain.

## Introduction

The microenvironment surrounding a brain injury and the cellular responses elicited in the remaining cells are key determinants of how efficiently a repair process will unfold. An important factor underlying the effectiveness of regenerative responses to an injury is the plasticity of the stem/progenitor cells in a tissue (Burda and Sofroniew, 2014). The degree to which the microenvironment and specific cell types within it provide pro- or anti-regenerative factors is highly context dependent. The neonatal mouse cerebellum has a remarkable capacity to regenerate cells ablated around birth (Wojcinski et al., 2017, Bayin N. S., 2021, Bayin et al., 2018, Altman and Anderson, 1971). Thus, the cerebellum provides an ideal system to study the roles that signals in the microenvironment play in key steps of the repair process in the brain.

The cerebellum is a folded hindbrain structure that is critical for motor coordination. It also participates in higher order social and cognitive behaviors through its circuit connections with all other brain regions (Badura et al., 2018, Buckner, 2013, Burda and Sofroniew, 2014, Salman and Tsai, 2016, Strick et al., 2009, Tomlinson et al., 2013). Compared to the rest of the brain, the cerebellum has protracted development, as its major growth occurs during the first two weeks after birth in mice and at least six months surrounding birth in humans (Altman and Bayer, 1997, Rakic and Sidman, 1970, Dobbing and Sands, 1973). This timing of the major growth of the cerebellum makes it susceptible to injury around birth. Indeed, cerebellar hypoplasia is the second leading risk factor for autism spectrum disorders and cerebellar injury around birth can have devastating outcomes and significant effects on subsequent quality of life (Tsai et al., 2018, Stoodley et al., 2017, Wang et al., 2014). Therefore, it is critical to better understand the regenerative processes that allow repair of the cerebellum.

All the cell types in the cerebellum are derived from two progenitor zones, the embryonic rhombic lip and the ventricular zone that give rise to the excitatory neurons, or the inhibitory neurons and glia, respectively (Leto et al., 2015, Joyner and Bayin, 2022). During postnatal growth, the rhombic lip-derived granule cell precursors (GCPs) cover the surface of the cerebellum in a structure named the external granule layer (EGL) and continue to proliferate in a sonic hedgehog (SHH) dependent manner for two weeks after birth in mice (Wechsler-Reya and Scott, 1999, McMahon et al., 2003, Corrales et al., 2006). Following their exit from the cell cycle, the granule cells (GC) migrate inwards to form the internal granule layer (IGL). Other SHH-dependent progenitor populations of the neonatal cerebellum are either gliogenic Nestin-Expressing Progenitors (NEPs) that express SOX2 and generate astroglia (astrocytes and Bergman glia) or neurogenic-NEPs that generate late born interneurons (Bayin N. S., 2021, Cerrato et al., 2018, Parmigiani et al., 2015). Gliogenic-NEPs reside either in the Bergmann glia layer (BgL) intermixed with Purkinje cells and generate Bergmann glia (Bg) and astrocytes, or in the white matter in the center of the lobules (folds) and generate astrocytes. Neurogenic-NEPs are restricted to the white matter and produce interneurons that migrate outwards to the outermost molecular layer (Bayin N. S., 2021, Brown et al., 2020, De Luca et al., 2015). Surprisingly, when the GCPs are killed upon injury soon after birth, the gliogenic-NEPs in the BgL (BgL-NEPs) undergo adaptive reprogramming to generate GCPs and replenish the EGL via a transitory cellular state that involves upregulation of the neurogenic gene *Ascl1* to promote a glial-to-neural fate switch (Wojcinski et al., 2017, Bayin N. S., 2021). Adaptive reprogramming involves multiple sequential stages starting with increased proliferation of BgL-NEPs, then a fate switch to neuronal progenitors, migration to the site of injury (EGL) and acquisition of a GCP identity. The full repertoire of injury-induced signals that initiate and govern adaptive reprogramming remains to be discovered.

In the adult brain, numerous cell types communicate and provide a concerted response to injury, including astrocytes, microglia (macrophages of the brain) and stem cells of the neurogenic niches (Frik et al., 2018). The timelines of the cellular responses of each cell type to injury - cell death, activation of microglia, reactive gliosis, proliferation, scar formation and cellular remodeling - have been delineated for specific adult brain injuries, particularly in the cerebral cortex. For example, upon traumatic brain injury cells release damage-associated molecular patterns (DAMPs), which act as an inflammatory stimulus and activate microglia that can lead to gliosis, eventually causing neurotoxicity and scarring (Donat et al., 2017). However, the cellular composition and microenvironment of the early postnatal brain are very different from the adult. In the neonatal cerebellum, microglia are immature (Li et al., 2019) and are still being generated, and NEPs and GCPs are actively proliferating and producing astroglia and neurons. Therefore, the existing knowledge on how adult brain cells react to injury might not apply to the neonatal cerebellum. For example, in the spinal cord, while neonatal microglia and astrocytes facilitate scarless repair, the same cells in the adult promote scarring upon spinal cord injury in mice (Li et al., 2020). It is thus important to study the microenvironment of the neonatal brain during repair to determine what factors promote or inhibit regeneration.

Dying cells release many factors, including reactive oxygen species (ROS) that activate signaling cascades in neighboring cells. However, little is known about how these signals regulate brain repair, especially during development. The level of ROS during homeostasis is regulated by metabolic processes, and typically is increased following injury (Niethammer, 2016). Furthermore, ROS can directly react with proteins that regulate proliferation, viability, quiescence or differentiation and metabolism (Bigarella et al., 2014, Tan and Suda, 2018). Thus, ROS are considered key signaling molecules that participate in the crosstalk between progenitor cell fate decisions and metabolic switches in a context- and cell type-dependent manner (Bigarella et al., 2014). One significant mechanism by which ROS signaling is implicated during inflammatory responses following an injury is through the activation of microglia, which in turn can lead to more ROS production (Smith et al., 2022). This process is critical as it can potentially promote repair. However, the role of ROS signaling during adaptive reprogramming of NEPs following neonatal cerebellar injury is not known.

Here we first delineate the sequential changes in the microenvironment upon injury (focused irradiation) to the mouse hindbrain at postnatal day 1 (P1). We then demonstrate a requirement for a transient increase in ROS levels at ∼24 hours (hr) post injury for cerebellar regeneration. Single cell RNA-sequencing (scRNA-seq) and bulk Assay for Transposase-Accessible Chromatin with sequencing (ATAC-seq) analyses revealed increased ROS signaling compared to controls that peaks 24 hr after injury in NEPs, demonstrating that ROS is an acute signal associated with the NEP response to GCP death. A functional role of ROS signaling was established using a transgene (*mCAT*) that expresses the human mitochondrial Catalase which can reduce ROS levels broadly. Several key steps in adaptive reprogramming were abrogated in *mCAT*/+ mice leading to reduced replenishment of the EGL and a smaller adult cerebellum. Finally, we show that the density of microglia is reduced at P5 in irradiated *mCAT* mice compared to controls and provide preliminary evidence that microglia play a role in the step of replenishing the EGL with BgL-derived GCPs during adaptive reprogramming.

## Materials and Methods

### Animals

All the mouse experiments were performed according to protocols approved by the Institutional Animal Care and Use Committee of Memorial Sloan Kettering Cancer Center (MSKCC) (protocol no. 07-01-001). Animals were housed on a 12-hour light/dark cycle and given access to food and water ad libitum.

Two mouse lines were used in this study: *Nes-Cfp* (JAX #034387) (Encinas et al., 2006) and *mCAT* (JAX #016197) (Schriner et al., 2005). Animals were maintained on an outbred Swiss Webster background. Both sexes were used for analyses except for the genomics experiments (scRNA-seq and ATAC-seq) where males were used.

#### EdU administration

5-ethynyl-2’-deoxyuridine (EdU) stock was dissolved in sterile phosphate-buffered saline (PBS) at 10 mg/mL and a dose of 5 μg/g was intraperitoneally injected into animals 1 hour prior to euthanasia.

#### PLX5622 administration

PLX5622 powder was provided by Plexxikon under a Materials Transfer Agreement. PLX5622 powder was first diluted in DMSO at 20mM and then diluted 1X in PBS just before intraperitoneal injection into newborn pups. Injections were given every day from P1 to P8 at a dose of 10µg/g of body weight. Control pups were injected with PBS-DMSO vehicle control.

#### Irradiation

P1 mice were anesthetized by hypothermia and given a single dose of ∼5Gy γ-irradiation in an X-RAD 225Cx (Precision X-ray) Microirradiator in the MSKCC Small-Animal Imaging Core facility. A 5-mm diameter collimator was used to target the hindbrain from the left side of the animal.

### Tissue preparation and histology

For immunocytochemistry, animals younger than P8 were sacrificed and then brains were dissected, fixed in 4% paraformaldehyde for 24 hr at 4°C, cryoprotected in 30% sucrose in phosphate-buffered saline (PBS) until they sank and then frozen in Cryo-OCT (Tissue-Tek). Older animals were anesthetized and then perfused with cold PBS followed by 4% paraformaldehyde prior to brain dissection. Frozen brains were cryosectioned sagittally at 14 μm and slides stored at -20°C. Midline cerebellar sections were used for quantification in all downstream analyses.

For immunofluorescence staining, slides were allowed to warm to room temperature (RT) and washed 3 times in PBS. Then, tissues were blocked for one hour with blocking buffer (5% BSA (w/v) in 1XPBS with 0.1% Triton-X) at RT. Primary antibodies diluted in the blocking buffer were placed on slides for overnight incubation at 4°C. Slides were then washed in PBS with 0.1% Triton-X and incubated with fluorophore-conjugated secondary antibodies diluted in the blocking buffer for 1 hr at RT. Following washes in PBS with 0.1% Triton-X after the secondary antibody incubation, nuclei were counterstained with Hoechst (1:3000) and the slides were mounted using mounting media (Electron Microscopy Sciences). Primary antibodies used are described in Table S1 and secondary antibodies were Alexa Fluor-conjugated secondary antibodies (1:1000).

EdU was detected using a Click-it EdU (Invitrogen, C10340) assay with Sulfo-Cyanine5 azide (Lumiprobe Corporation, A3330).

For TUNEL staining, after primary antibody incubation and washes, sections were permeabilized in PBS with 0.5% TritonX-100 for 10 minutes (min) and then pre-incubated in TdT buffer (30mM Tris HCl, 140 mm sodium cacodylate and 1mM CoCl_2_) for 15 min at RT. Slides were then incubated for 1hr at 37°C in TUNEL reaction solution containing Terminal Transferase and Digoxigenin-11dUTP (Roche). After the TUNEL reaction and washes, slides were incubated with a secondary antibody solution which included a sheep anti-dixogenin-rhodamine (Roche) for the visualization of TUNEL reaction.

### Image acquisition and analysis

Images were collected with a DM6000 Leica microscope, a NanoZoomer Digital Pathology microscope (Hamamatsu Photonics), or an LSM880 confocal microscope (Zeiss). Images were processed using NDP.view2 software, ImageJ software (NIH, Bethesda, MA, USA) and Photoshop (Adobe).

### Cell dissociation for FACS and flow cytometry

Cerebella were dissected into ice-cold 1x Hank’s Buffered Salt Solution (Gibco) and dissociated using Accutase (Innovative Cell Technologies) at 37°C for 10-15 min. After dissociation, Accutase was diluted with 3x excess volume of neural stem cell media (Neurobasal medium, supplemented with N2, B27 (without vitamin A)), and nonessential amino acids (Life Technologies, Gibco) and cells were filtered using a 40 μm mesh cell strainer. After 5 min of centrifugation in a chilled centrifuge at 500g, the pellet was resuspended in neural stem cell media and strained using strainer cap tubes (Falcon) for downstream experimentation. All centrifugation was performed at 4°C and cells were kept on ice when possible.

### Flow cytometry analysis for ROS and mitochondria mass

For MitoSOX and MitoTracker analyses, cells were incubated for 30 min at 4°C with 5 μM of MitoSOX and 100 μM of MitoTracker (Thermo Fisher Scientific) to assess mitochondrial superoxide production and mitochondrial mass, respectively. Data were collected using a LSR Fortessa flow cytometer (BD) and analyzed using FlowJo software. The gating for the MitoSOX or MitoTracker high populations (top 90%) were performed as previously described (Clutton et al., 2019)

### Multiplexed sc-RNAseq

#### Sample preparation

2-4 *Nes-Cfp/+* cerebella/replicates from male control non-irradiated pups or pups that were irradiated at P1 were collected at P1 (control only), P2, P3 and P5 and dissociated as described above. All conditions were performed in 2 replicates for nonIR and IR conditions, except for P5. Following dissociation, CFP+ cells were immediately sorted on a BD FACS Aria sorter (BD Biosciences) using a 100 μm nozzle. 50,000 Nes-CFP+ cells from each sample were processed for scRNA-seq. Cells were sorted into neural stem cell media.

#### Multiplexing, droplet preparation, sequencing and data processing

The scRNA-Seq of FACS-sorted Nes-CFP+ cell suspensions was performed on a Chromium instrument (10X genomics) following the user guide manual for 3′ v3.1. In brief, 50,000 FACS-sorted cells from each condition were multiplexed using CellPlex reagents (10x Genomics) as described by the manufacturer’s protocol. P3 nonIR replicate #2 did not yield sufficient cells after multiplexing. The viability of cells was above 95%, as confirmed with 0.2% (w/v) Trypan Blue staining and barcoded cells were pooled into a single sample in PBS containing 1% bovine serum albumin (BSA) to a final concentration of 700–1,300 cells per μl. 2-3,000 cells were targeted for each sample. Samples were multiplexed together on one lane of a 10X Chromium following the 10x Genomics 3’ CellPlex Multiplexing protocol and a total of ∼30,000 cells were targeted for droplet formation. Cells were captured in droplets. After the reverse transcription and cell barcoding in droplets, emulsions were broken, and cDNA was purified using Dynabeads MyOne SILANE followed by PCR amplification per the manufacturer’s instruction. Final libraries were sequenced on an Illumina NovaSeq S4 platform (R1 – 28 cycles, i7 – 8 cycles, R2 – 90 cycles) by the MSKCC core facility.

#### Data Analysis

Single-cell RNA-seq fastq files were demultiplexed using sharp (https://github.com/hisplan/sharp) and initially mapped to the mouse mm10 reference genome using Cell Ranger v6.0.1 (Zheng et al., 2017). The Cell Ranger BAM files for individual samples were then converted back to fastq files using *bamtofastq* (Cell Ranger v7.0.1), with --reads-per-fastq=1000000000. Reads were then mapped to the GRCm39 mouse reference with Gencode annotations (vM30) using STARsolo v2.7.10a (--soloFeatures Gene Velocyto, --soloType CB_UMI_Simple, --soloCBwhitelist 3M-february-2018.txt, --soloUMIlen 12, --soloCellFilter EmptyDrops_CR, --soloMultiMappers EM) (Kaminow et al., 2021, Frankish et al., 2021).

The STARsolo output was used to create Seurat (v4.3) objects for each sample with spliced and unspliced read count matrices (Hao et al., 2021). The objects were integrated, by running NormalizeData and FindVariableFeatures (selection.method = “vst”, nfeatures = 3000) on each one, then SelectIntegrationFeatures and FindIntegrationAnchors on the list of objects and finally IntegrateData with the anchors. Counts were then normalized with SCTransform. Based on manual analysis, cells were filtered out where number of detected genes was ≤ 1,500, number of detected transcripts was ≥ 40,000 and mitochondrial gene percentage ≥ 5%. To determine cell cycle phases, the Kowalczyk et al. (Kowalczyk et al., 2015) cell cycle markers were used, assuming the gene names, capitalized to the title case, to be orthologous between mouse and human with the CellCycleScoring function. SCTransform was used to regress out the “Cell Cycle difference” score (S score – G2M score). Dimension reduction was performed using RunPCA and RunUMAP (dims = 1:20, n.neighbors = 30). For clustering, FindNeighbors was run using the first 20 principal components, then FindClusters with the Leiden algorithm with the default settings (Traag et al., 2019). Clusters were annotated using known markers: GCPs (*Atoh1+*/*Barhl1+*/*Tubb3+*), BgL-NEPs (*Hopx+*), ependymal cells (*Foxj1+*), immature interneurons (*Pax2+*), Neurogenic NEPs (*Ascl1+*), astrocytes (*Slc6a11+*), oligodendrocytes (*Olig1+*), meninges (*Col3a1+*/*Vtn+*/*Dcn+*), microglia (*Ly86+*/*Fcer1g+*).

The differential gene expression analyses were performed individually on *Hopx+* gliogenic-NEPs, *Ascl1+* neurogenic-NEPs, and GCPs following the same computational workflow. First, clusters containing *Hopx*-NEPs (clusters 2, 3, 6, 10), *Ascl1*-NEPs (clusters 5, 8, 11), or GCPs (clusters 1, 4, 7, 12, 14) were subsetted from the original data set based on the expression of *Hopx*, *Ascl1*, and *Atoh1*, *Barhl1* and *Rbfox*, respectively. Second, the subsetted cells were divided according to whether the cells were from P2 or P3+P5 and split based on their biological replicate. The split data sets were normalized using NormalizeData with default parameters, and the 3,000 top variable features were computed using FindVariableFeatures with default settings. Re-integration of the data sets was subsequently performed using SelectIntegrationFeatures, FindIntegrationAnchors, and IntegrateData all with default parameters as previously described, except for IntegrateData in the *Ascl1+* neurogenic-NEP analyses where k.weight = 50 was used. Following re-integration, SCTransform with default parameters was used to normalize mitochondrial read percentage, cell cycle difference score, number of features, and number of counts. Dimension reduction was thereafter performed using RunPCA with default parameters and RunUMAP with default parameters except dims = 1:40, repulsion.strength = 0.1, and min.dist = 0.5. Clustering was subsequently performed using FindNeighbors with default settings except dim = 30 and FindClusters with default settings for resolutions between 0.1 and 3 using the original Louvain algorithm. A final resolution which gave a high silhouette score with a relatively low negative silhouette proportion was selected for individual data sets. To allow downstream DESeq2 analyses, count matrices were constructed by aggregating counts from cells from the same treatment condition and biological replicate using AggregateExpression. The aggregated count matrices were converted into a DESeq2 dataset object using DESeqDataSetFromMatrix, grouping the samples by treatment conditions. Genes with fewer than 10 counts were filtered out. DESeq2 (v1.36.0) was used to perform the differential expression analyses using a negative binomial distribution and default settings (Love et al., 2014). The results were visualized using EnhancedVolcano with a fold change cut-off of ±0.5 and an adjusted p-value threshold of 0.05.

The GO term analyses were performed on differentially expressed genes from the DESeq2 results filtered with a log2 fold change threshold of ±0.5 and an adjusted p-value threshold of 0.05 for each comparison. The probability weight function was computed using nullp with default parameters and the mm8 mouse genome. The background genes used to compute the GO term enrichment includes all genes with gene symbol annotations within mm8. The category enrichment testing was performed using goseq with default parameters from the goseq package (v1.48.0).

### Bulk ATAC-seq

#### Sample preparation

FACS-sorted Nes-CFP+ cells (30,000-50,000 per replicate) were isolated from control or irradiated (at P1) P2 cerebella. 2-3 cerebella were pooled for each sample. Cells were immediately processed for nuclei preparation and transposition using the OMNI-ATAC protocol (Corces et al., 2017). Sequencing was performed at the MSKCC genomics core facility using the Illumina NovaSeq S4 platform.

#### Data Processing and Analysis

Raw sequencing reads were 3’ trimmed and filtered for quality and adapter content using version 0.4.5 of TrimGalore (https://www.bioinformatics.babraham.ac.uk/projects/trim_galore), with a quality setting of 15, and running version 1.15 of cutadapt and version 0.11.5 of FastQC. Version 2.3.4.1 of bowtie2 (http://bowtie-bio.sourceforge.net/bowtie2/index.shtml) was used to align reads to mouse assembly mm10 and alignments were deduplicated using MarkDuplicates in version 2.16.0 of Picard Tools. Enriched regions were discovered using MACS2 (https://github.com/taoliu/MACS) with a p-value setting of 0.001, filtered for blacklisted regions (http://mitra.stanford.edu/kundaje/akundaje/release/blacklists/mm10-mouse/mm10.blacklist.bed.gz), and a peak atlas was created using +/- 250 bp around peak summits. The BEDTools suite (http://bedtools.readthedocs.io) was used to create normalized bigwig files. Version 1.6.1 of featureCounts (http://subread.sourceforge.net) was used to build a raw counts matrix and DESeq2 was used to calculate differential enrichment for all pairwise contrasts. Peak-gene associations were created by assigning all intragenic peaks to that gene, while intergenic peaks were assigned using linear genomic distance to transcription start site. Network analysis was performed using the assigned genes to differential peaks by running enrichplot::cnetplot in R with default parameters. Composite and tornado plots were created using deepTools v3.3.0 by running computeMatrix and plotHeatmap on normalized bigwigs with average signal sampled in 25 bp windows and flanking region defined by the surrounding 2 kb. Motif signatures were obtained using Homer v4.5 (http://homer.ucsd.edu) on differentially enriched peak *regions*.

### Data and code availability

The scRNA-seq data was submitted to ArrayExpress (Accession E-MTAB-13353). Bulk ATAC-seq data has been submitted to GEO (GSE269342). The code used to carry out the scRNA-seq analysis can be found on GitHub repository: https://github.com/BayinLab/Pakula_et_al_23

### Quantification and statistical analysis

For detecting TUNEL, IBA1, GFAP, CFP and SOX2, images were captured using a DM600 Leica fluorescent microscope and subsequently quantified on ImageJ Software (NIH). Measurements were conducted on lobules 3, 4 and 5 of midsagittal sections unless indicated in Figure legends.

Positive cells were counted and densities were calculated based on BgL length, EGL area, WM area or on the whole cerebellum without the EGL (outside EGL) as indicated in the figures. For the cerebellar section area, images were acquired using a Nanozoomer2.0 HT slide scanner (Hamamatsu). Midsagittal sections were selected and exported for manual analysis using ImageJ software. For EGL thickness, images of lobules 3, 4 and 5 from midsagittal sections were obtained using a DM600 Leica fluorescent microscope and analyzed on ImageJ. EGL thickness was calculated as the EGL area divided by the EGL perimeter.

Prism (GraphPad) was used for all statistical analyses. Statistical tests performed in this study were Welch’s two-tailed t-test and Two-way analysis of variance (ANOVA) followed by post hoc analysis with Tukey’s multiple comparison tests. A p-value ≤ 0.05 was considered as significant. Results are presented as the mean ± SEM. At least three biological samples and 2 to 3 sections per sample were analyzed for each experiment to ensure reproducibility and the sample sizes are reported in the Results section for significant data. For qualitative analysis, midsagittal sections from at least 4 samples were observed.

## Results

### scRNA-seq of NEPs during adaptive reprogramming reveals increased cellular stress, ROS signaling and DNA damage

To investigate the molecular changes that NEP subpopulations undergo upon injury to the EGL, in particular an increase in the signaling pathways associated with injury induced cellular stress and ROS, we performed multiplexed scRNA-seq of CFP+ cells isolated by fluorescence-activated cell sorting (FACS) of cerebella from *Nes-Cfp/+* transgenic neonates either irradiated at P1 (IR; P2, P3, P5) or non-irradiated (nonIR; P1, P2, P3, P5) (Figure 1A, Supplementary Figure 1A, B). Following the filtering out of poor quality cells and integration of replicates and the two conditions, the clustering of 11,878 cells (6,978 nonIR and 4,900 IR) was performed (Hao et al., 2021). The analysis revealed the expected three distinct groups of cells: gliogenic-NEPs and astrocytes (*Hopx*+ clusters 2, 3, 6, 10), neurogenic-NEPs (*Ascl1*+ clusters 5, 8, 11) and GCP (*Atoh1*+ clusters 1, 4, 7, 12, 14) that were present at each stage and in both conditions (Figure 1B-E, Supplementary Figure 1C-F, Table S2). These groups of cells were further subdivided into molecularly distinct clusters based on marker genes and their cell cycle profiles or developmental stages (Figure 1D, Table S2). In addition, oligodendrocyte progenitors (cluster 15), microglia (clusters 17, 21), ependymal cells (clusters 13, 18, 19) and meninges (cluster 16) were detected (Figure 1B, D, Table S2). Cluster 20 represented low-quality cells and was omitted from downstream analyses. As expected, the GCP clusters were enriched in the cells from irradiated mice and at P5 (Supplementary Figure 1C). Detection of *Nes* mRNA confirmed that the transgene reflects endogenous *Nes* expression in progenitors of many lineages, and also that the perdurance of CFP protein in immediate progeny of *Nes*-expressing cells allowed the isolation of these cells by FACS (Figure 1E).

**Figure 1:**
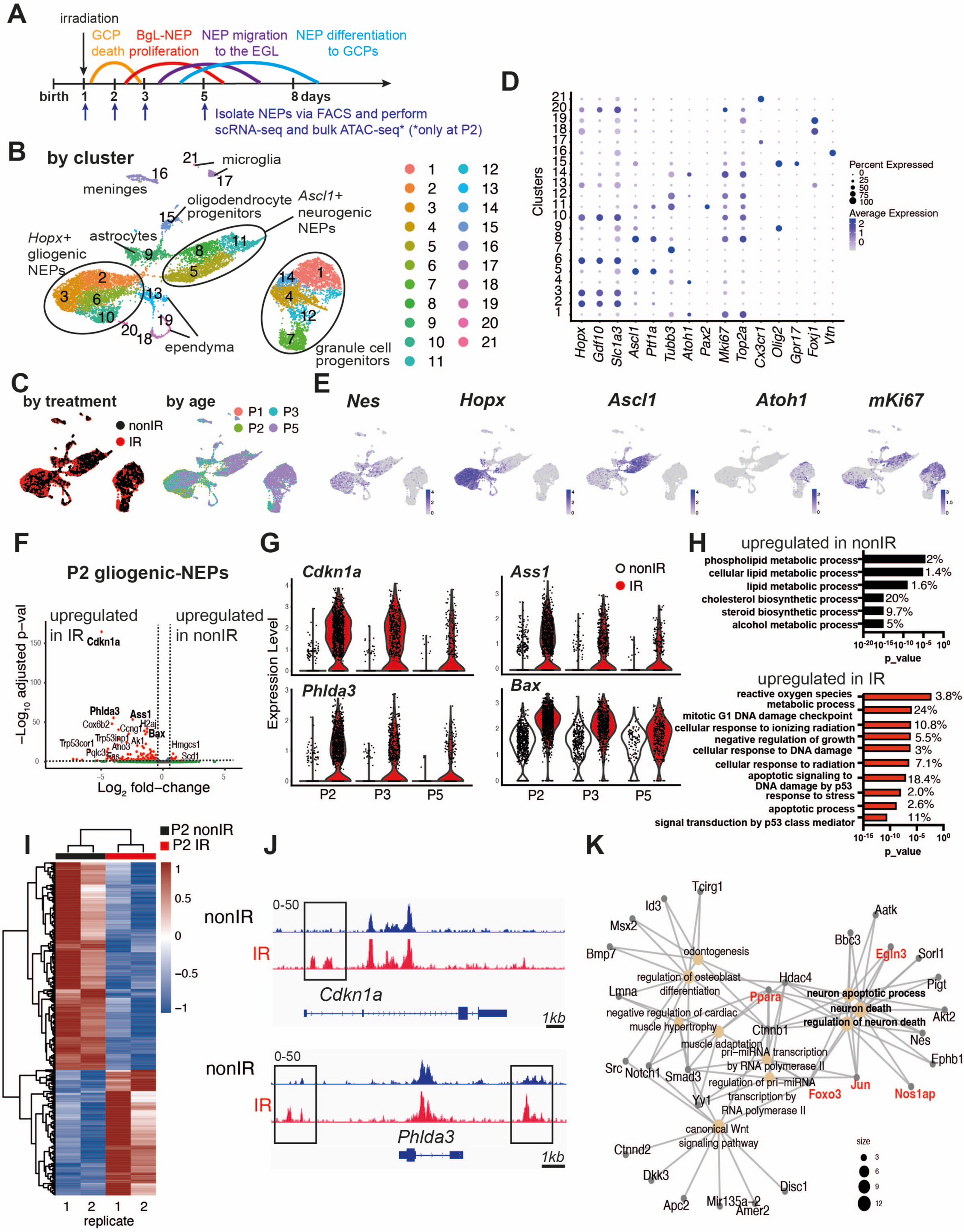
Injury induces ROS and cell stress signaling reflected by changes in the transcriptome and chromatin landscape of progenitors. **(A)** Schematic summarizing the experimental plan. **(B-C)** UMAP projections of 11,878 cells (6,978 nonIR and 4,900 IR) showing cluster annotations (B), treatment (black: nonIR, red: IR) and the age of the samples (red: P1, green: P2, blue: P3, purple: P5) (C). **(D)** Dot plot showing the expression levels of key marker genes used for cluster annotation (gliogenic-NEPs: *Hopx, Gdf10, Slc1a3*, neurogenic-NEPs: *Ascl1, Ptf1a*, immature neurons: *Pax2*, GCPs: *Atoh1*, postmitotic neurons: *Tubb3*, microglia: *Cx3cr1*, oligodendrocyte progenitors: *Olig2*, oligodendrocytes: *Gpr17*, Ependymal cells: *Foxj1*). **(E)** Feature plots showing *Nes*, *Hopx* (gliogenic-NEPs), *Ascl1* (neurogenic-NEPs) and *Atoh1* (GCPs) and *mKi67* (proliferation) expression highlighting the three main populations of interest. Clusters containing *Hopx*-NEPs (clusters 2, 3, 6, 10), *Ascl1*-NEPs (clusters 5, 8, 11), or GCPs (clusters 1, 4, 7, 12, 14) were subsetted from the original data set and were divided according to age (P2 or P3+P5) for the downstream differential expression analyses. **(F)** Volcano plot showing differentially expressed genes in the P2 gliogenic-NEPs (red: adjusted p-value≤0.05, log_2_fold-change>|1|). **(G)** Violin plots showing some of the top differentially expressed genes in P2 gliogenic-NEPs and how their expression changes over time with respect to their expression in control cells. **(H)** Some of the significant GO terms associated with differentially expressed genes in P2 gliogenic-NEPs that were either upregulated in nonIR (top panel) or IR (bottom panel) cells (adjusted p-value≤0.05, Table S3). **(I)** Heatmap showing differentially open chromatin regions in P2 nonIR and IR NEPs, identified by bulk ATAC-seq (1168 differentially open regions, adjusted p-value<0.05, Table S4). **(J)** Tracks highlighting the injury-induced open chromatin regions around *Cdkn1a* and *Phlda3,* the top differentially expressed genes identified in (F). **(K)** Linkages between genes and GO-terms identified by the ATAC-seq data revealed an active transcriptional network involved in regulating cell death and apoptosis. Genes colored in red (*Ppara*, *Egln3*, *Foxo3*, *Jun* and *Nos1ap)* have been implicated as upregulated with increased ROS levels or involved in ROS signaling.

Our further analyses focused on changes in the signaling pathways associated with injury induced cellular stress and ROS. We previously showed that a subset of the *Hopx+* gliogenic-NEPs that are present in the BgL are the ones that undergo adaptive reprograming following GCP death (Wojcinski et al., 2017, Bayin N. S., 2021). We therefore assessed the immediate and later effects of GCP death on *Hopx+* gliogenic-NEPs by performing differential expression analyses between nonIR and IR gliogenic-NEPs (*Hopx+,* clusters 2, 3, 6, 10) at P2, or at P3 and P5 (P3+5). 24 hr after injury at P1, 132 genes in gliogenic-NEP clusters were significantly upregulated in IR compared to 34 genes that were upregulated in nonIR P2 cells (adjusted p-value≤0.05, Figure 1F, Table S3). The significantly increased genes included *Cdkn1a, Phlda3, Ass1* and *Bax,* all of which have been implicated as increased in response to DNA damage and in ROS signaling, as well as in anti-apoptotic functions (Figure 1G) (Masgras et al., 2012, Bensellam et al., 2019, Qiu et al., 2014, Jiang et al., 2008). Indeed, many of the significant gene ontology (GO) terms associated with the genes upregulated in gliogenic-NEPs with injury were related to response to irradiation, DNA damage, the P53 pathway and ROS metabolic processes, whereas many of the significant GO terms associated with the genes upregulated in the nonIR cells at P2 were related to metabolic processes (p-value≤0.05, Figure 1H, Table S4). Interestingly, the transcriptional changes observed at P2 were less pronounced at later time points, where although some of the top differentially expressed genes at P2 were still significantly upregulated at P3+5 stages combined (e.g. *Cdkn1a, Phlda3, Ass1, Bax*), the increase in expression levels of these genes upon injury and/or the number of cells expressing them gradually declined after P2 in IR gliogenic-NEPs when compared to their nonIR counterparts (Figure 1G). Genes upregulated in IR P3+5 gliogenic NEPs were associated with similar GO terms to those at P2, such as response to irradiation and the P53 pathway, however, “ROS metabolic processes” was no longer a significantly enriched GO term (p-value≤0.05, Supplementary Figure 2A, F, Table S3-4).

To further assess the injury induced transcriptional signatures, we performed the same differential expression analysis on nonIR and IR neurogenic-NEPs (*Ascl1+,* clusters 5, 8, 11) and GCPs (*Atoh1+,* clusters 1,4,7,12,14) at P2, or at P3+P5 to identify the immediate and later changes upon injury at birth. Some of the DNA damage and apoptosis related genes that were upregulated in IR gliogenic-NEPs (*Cdkn1a*, *Phlda3*, *Bax*) were also upregulated in the IR neurogenic-NEPs and GCPs at P2 (Supplementary Figure 2B-E). Similar to the gliogenic-NEPs, P2 IR neurogenic-NEPs showed significant upregulation of genes associated with GO terms related to stress response and apoptosis following injury, although the ROS related GO term was not as prominent (Supplementary Figure 2B, G, Table S3-4). Interestingly, although IR neurogenic-NEPs at P3+5 had only 27 genes that were significantly upregulated following injury (adjusted p-value≤0.05, Table S3, Supplementary Figure 2C), the genes included ones associated with neural stem cells and BgL-NEPs (e.g. *Id1, Apoe, Ednrb*). The latter genes could represent the transitory *Ascl1+* BgL-NEP population that induces *Ascl1* expression during adaptive reprogramming and would be included in the *Ascl1+* neurogenic clusters or reflect the delayed neurogenesis of neurogenic-NEPs previously demonstrated (Bayin N. S., 2021). Consistent with the latter, the nonIR P3+5 neurogenic-NEPs showed an increase in mature neuron markers compared to IR cells (adjusted p-value<0.05, Supplementary Figure 2C, H, Tables S3-4).

In contrast to the gliogenic- and neurogenic-NEP subtypes, P2 IR GCPs showed upregulation of genes enriched in GO terms such as neural differentiation and axonogenesis as well as ROS signaling and nonIR P2 GCPs showed increased expression of genes involved in cell cycle and mitosis (Supplementary Figure 2D, I). This result could reflect the death of highly proliferative GCPs after irradiation and sparing of only postmitotic granule cells upon irradiation at P1. P3+5 IR GCPs showed increased expression of genes associated with BgL-NEPs (*Id1* and *Gdf10*, adjusted p-value ≤ 0.05) and genes associated with GO terms such as cell cycle and cell fate commitment whereas P3+5 nonIR GCPs showed enrichment for GO terms related to cell migration and neurogenesis (adjusted p-value ≤0.05, Supplementary Figure 2D, E, I, J, Tables S3-4). This result could reflect the delayed neurogenesis of GCPs following injury. Interestingly, we did not observe significant enrichment for GO terms associated with cellular stress response in the GCPs that survived the irradiation compared to controls, despite significant enrichment for ROS signaling related GO-terms (Table S4). Collectively, these results indicate that injury induces significant and overlapping transcriptional changes in NEPs and GCPs. The gliogenic- and neurogenic-NEP subtypes transiently upregulate stress response genes upon GCP death, and an overall increase in ROS signaling is observed in the injured cerebella.

### Injury induces changes in NEP chromatin landscape at P2

We next tested whether GCP death at birth induces changes to the chromatin landscape of NEPs that reflect the altered gene expression observed with scRNA-seq, by performing bulk ATAC-seq on FACS-isolated CFP+ cells from P2 control and injured *Nes-Cfp/+* pup cerebella (Figure 1A). P2 was chosen as it is the stage when GCPs contribute the least to the total Nes-CFP+ FACS population and to identify the immediate effects of the injury on NEP chromatin. Analysis of differentially open chromatin showed that injury induces major changes to the chromatin landscape of the NEPs (1168 differentially open regions, adjusted p-value<0.05, Figure 1I. Table S5). Of interest, *Cdkn1a* and *Phlda3*, two genes stimulated by ROS and injury (Bensellam et al., 2019, Masgras et al., 2012) and that were upregulated in gliogenic-NEPs after irradiation (Figure 1F-G, Supplementary Figure 2B) exhibited new accessible regions around their gene bodies compared to the nonIR P2 Nes-CFP+ cells (Figure 1J). However, not all genes in the accessible areas were differentially expressed in the scRNA-seq data. While some of this could be due to the detection limits of scRNA-seq, further analysis is required to assess the mechanisms of how the differentially accessible chromatin affects transcription. Known motif analysis in the regions with increased accessibility upon injury showed enrichment for binding motifs of the JUN/AP1 transcriptional complex (p-value = 10^-14^, % of target sequences with motif = 15%, Table S6) which is known to act in response to cellular stress and be activated by ROS. In addition, the DNA binding site for FOXO3, a transcription factor that regulates ROS levels, had increased chromatin accessibility (p-value = 0.001, % of target sequences with motif = 22.41%) (Table S6) (Filosto et al., 2003, Auten and Davis, 2009, Hagenbuchner et al., 2012). Finally, linkages between the genes in differentially open regions identified by ATAC-seq and the associated GO-terms revealed an active transcriptional network involved in regulating cell death and apoptosis (Figure 1K). Furthermore, some of the genes involved in this response, such as *Ppara*, *Egln3*, *Foxo3*, *Jun* and *Nos1ap,* have been implicated as upregulated with increased ROS levels or involved in ROS signaling (Devchand et al., 2004, Kaelin, 2005, Hagenbuchner et al., 2012, Filosto et al., 2003). In summary, our ATAC-seq data analysis along with the scRNA-seq provide strong evidence that irradiation causes increased ROS signaling in the BgL-NEPs upon GCP death by inducing transcriptional and epigenetic changes within 1 day after injury (P2). In addition, genes related to cell survival and death, cellular stress and DNA damage are upregulated in NEPs shortly upon injury, possibly as a means to overcome the cellular effects of injury and induce adaptive reprogramming of NEPs to replenish the lost cells.

### Transient increase in cerebellar ROS during apoptosis of granule cell precursors

To validate that the transient increase in expression of genes associated with cellular stress and ROS signaling is due to an increase in ROS upon cerebellar injury, we quantified ROS levels in whole cerebellum samples using a mitochondrial superoxide indicator (MitoSOX) via flow cytometry. A significant increase in ROS (cells present in the top 90% MitoSOX+ intensity) was observed specifically at P2 (p=0.0005, n≥10) and not at P3 or P5 in IR pups compared to nonIR (Figure 2A, B and Supplementary Figure 3A, B). Furthermore, quantification of mitochondrial mass with MitoTracker revealed a reduction only at P2 (p=0.0005, n≥10) in IR pups (Figure 2C and Supplementary Figure 3C-E). Additionally, quantification of TUNEL staining in midline sagittal sections of the cerebella showed a large increase in cell death in the EGL of injured cerebella at P2 (p=0.0040, n=4), but not at P3 compared to the controls (Figure 2D, E; see also Figure 3K, L). Outside the EGL there was a small increase in cell death after injury that was only significant at P3 (p=0.0379, n≥3) (Figure 2F, G). Thus, a transient increase in ROS in the cerebellum one day after irradiation at P1 correlates with the timing of the major death of GCPs.

**Figure 2:**
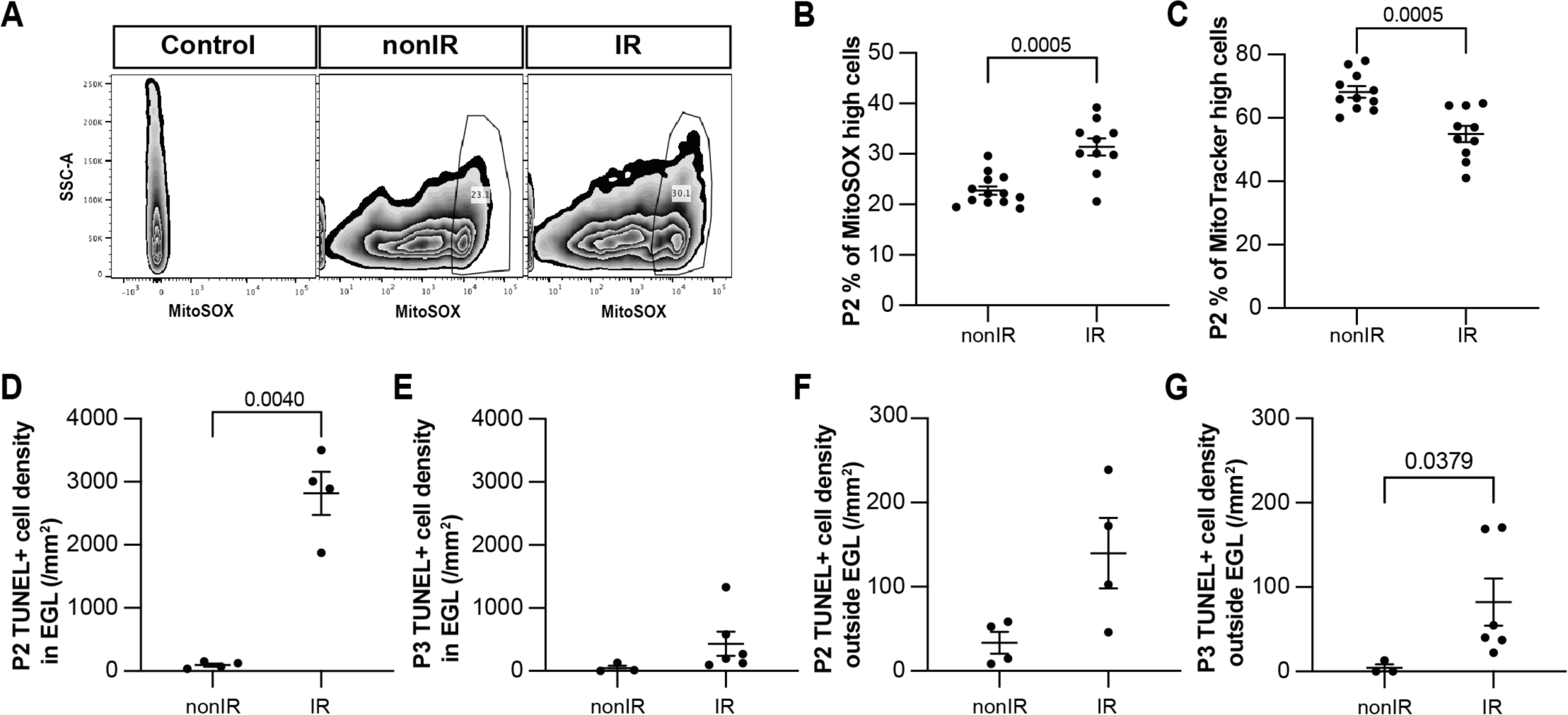
Cerebellar injury at P1 results in increased superoxide production, a reduction in mitochondria and increased cell death in the EGL that peaks 24h after injury. **(A)** Examples of flow cytometry analysis of mitochondrial ROS at P2 from nonIR and IR cerebella using MitoSOX dye. Gating determined the top 90% MitoSOX signal (MitoSOX high cells). **(B, C)** Quantification of MitoSOX high (B) and MitoTtracker high (C) expression in nonIR and IR cerebella at P2. **(D, E)** Quantification of TUNEL+ cell density in the EGL at P2 (D) and P3 (E) in lobules 3-5 of nonIR and IR mice. **(F, G)** Quantification of TUNEL+ cell density outside the EGL at P2 (F) and P3 (G) in lobules 3-5 of nonIR and IR mice. EGL, External granular layer; SSC, side scatter; P, postnatal day; nonIR, non-irradiated; IR, irradiated. All statistical significance was determined using an unpaired t-test and data are represented as mean ± SEM.

**Figure 3:**
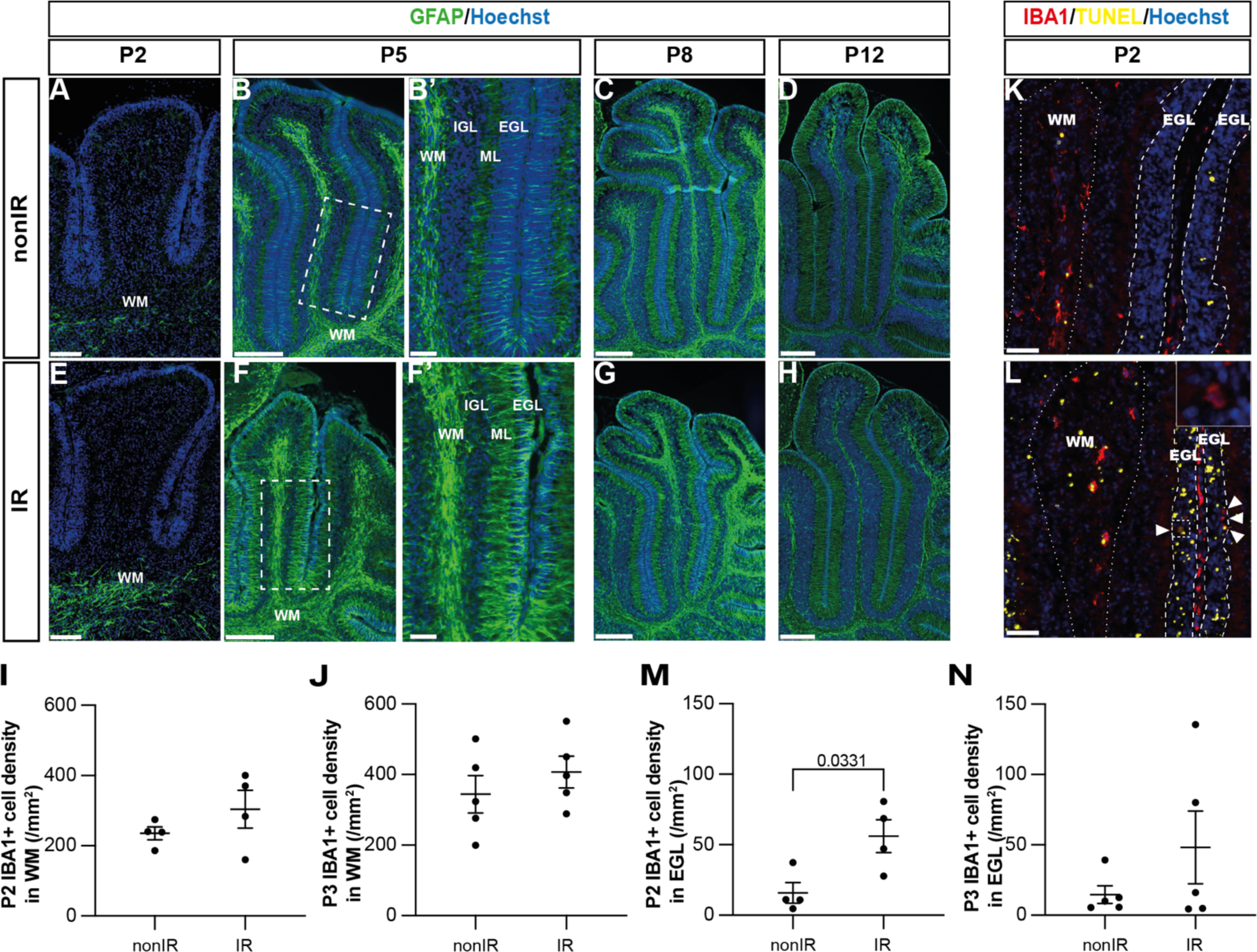
Cerebellar injury at P1 induces transient microglial recruitment to the EGL and prolonged astroglial microenvironment changes in the cerebellum. **(A-H)** Immunohistochemical (IHC) staining of medial sagittal cerebellar sections for GFAP (green) in lobules 4/5 of nonIR and IR cerebellum at the stages indicated. Nuclei were counterstained with Hoechst. (B’) and (F’) show high-power images of white dashed line boxes in (B) and (F), respectively. **(I, J)** Quantification of IBA1+ cell density in the WM at P2 (I) and P3 (J) in lobules 3-5 of nonIR and IR mice. **(K, L)** IHC staining of medial sagittal cerebellar sections for IBA1 and TUNEL in lobule 3 of nonIR and IR cerebellum at P2. Nuclei were counterstained with Hoechst. White matter (WM) and external granular layer (EGL) are delineated by white dotted lines and dashed lines, respectively. High-power image in (L) of the area indicated by the white dashed line represents an IBA1+ cell present in the EGL. White arrowheads indicate additional IBA1+ cells in the EGL. **(M, N)** Quantification of IBA1+ cell density in the EGL at P2 (M) and P3 (N) in lobules 3-5 of nonIR and IR mice. EGL, External granular layer; WM, White matter; P, postnatal day; nonIR, non-irradiated; IR, irradiated. Scale bar: A and E 100µm, B, C, D, E, F, G and H: 250µm, B’ and F’: 50µm, I and J: 50µm. All statistical significance was determined using an unpaired t-test and data are represented as mean ± SEM.

### Altered glial microenvironment following death of granule cell precursors

Given the potential importance of glial cells to regenerative cellular responses after injury, we next asked whether the glial microenvironment of the cerebellum changes during early postnatal development in response to irradiation at P1. We first analyzed the astrocyte marker GFAP, since it is generally upregulated soon after brain injury (Burda and Sofroniew, 2014). Most astrocytes, including the specialized Bg, express GFAP in the adult cerebellum, but the cells are generated by gliogenic-NEPs during the first two weeks after birth in mice. We therefore determined the normal location and timing of initiation of GFAP expression in these cells during postnatal cerebellum development. GFAP was first detected in rare astrocytes at P2 located in the white matter (WM) below the lobules, with strong expression in all WM astrocytes at P5 and later stages (P8 and P12) (Figure 3A-D, observed in n=4 mice/stage). In contrast, GFAP expression in astrocytes in the developing IGL and in the glial processes of Bg that project through the molecular layer (ML) and EGL was not detectable until P5 and was much stronger at P8 and P12 (Figure 3A-D). Interestingly, after injury at birth, GFAP expression was prematurely upregulated in the WM astrocytes below the lobules at P2 and in the remaining astrocytes at P5, including in the fibers of Bg that extend through the ML and EGL, compared to nonIR cerebella (Figure 3A-H, observed in n=4 mice/stage). GFAP expression was similar in all glia in both conditions at P8 and P12. These results reveal that astrocytes and Bg react to EGL injury caused by irradiation at P1 by initiating GFAP expression earlier than normal in the deep WM, and then the IGL and in Bg.

The macrophages of the brain, microglia, are generated in the early embryo but their main increase in cell number occurs during neonatal development in mice (Hammond et al., 2018). We found that nearly all microglia were located in the WM of the cerebellum, and that the density of IBA1+ microglia in the WM increased between P2 (235.1 ± 18.3 cells/mm^2^) and P3 (344.3 ± 53.1 cells/mm^2^) and then was maintained at P5 (350.4 ± 40.2 cells/mm^2^) (Fig. 3I-K, Supplementary Figure 3F). Since the area of the cerebellum is increasing between P2 and P5, active microglia production and/or infiltration must continue to occur between P2 and P5. As expected, after irradiation the density of microglia in the injured EGL was significantly increased (∼3 fold) at P2 but not at P3 or P5 compared to controls (p=0.0331, n=4) (Fig. 3L-N, Supplementary Figure 3G). No difference in the microglial density in the WM was detected between conditions at both time points (Fig. 3I-L; Supplementary Figure 3F). Thus, as expected microglia density was increased in the EGL one day after injury when the maximum GCP cell death occurs.

### Decreasing ROS impairs cerebellar repair

Given the transient increase in ROS signaling in gliogenic-NEPs during peak GCP death and recruitment of microglia to the EGL, and the later astroglial response, we tested whether an increase in ROS is necessary for adaptive reprogramming and cerebellar repair following EGL injury at P1. To reduce the level of ROS we utilized an *mCAT* transgenic mouse line that expresses the human mitochondrial catalase ubiquitously from a *CMV* promoter (Schriner et al., 2005). The transgene is expected to reduce mitochondrial ROS levels in all cells by catalyzing the breakdown of hydrogen peroxide into water and oxygen, hence protecting the cells from oxidative damage. We confirmed that mCAT protein is expressed throughout the cerebellum using immunohistochemical (IHC) staining of cerebellar sections (Figure 4A, B). MitoSOX flow cytometry revealed a significant decrease in ROS in the cerebella of *mCAT/+* mice 1 day after irradiation (P2) (p=0.0163, n≥8) and not at P3 or P5 compared to control IR mice and no baseline decrease in ROS at any stage in nonIR mice (Figure 4C and Supplementary Figure 4A, B). Mitochondrial mass, as measured by MitoTracker flow cytometry, was reduced at P2 in *mCAT/+* IR compared to nonIR *mCAT/+* pups with no significant difference observed between *mCAT/+* and control IR mice (Fig. 4D, Supplementary Figure 4C, D). The level of cell death in the EGL and density of IBA1+ microglia in the EGL and elsewhere in the cerebellum were similar between *mCAT/+* and littermate control IR mice at P2 and P3 (Fig. 4E, F, Supplementary Figure 4G, J). There was a slight increase in cell death outside the EGL at P2 in IR *mCAT/+* compared to nonIR *mCAT/+* mice but not compared to IR controls (Supplementary Figure 4F, E). Thus, the *mCAT* transgene counteracts the transient increase in cerebellar ROS following EGL injury but does have a major effect on GCP death or the infiltration of microglia to the injured EGL (Supplementary Figure 4 F-J).

**Figure 4:**
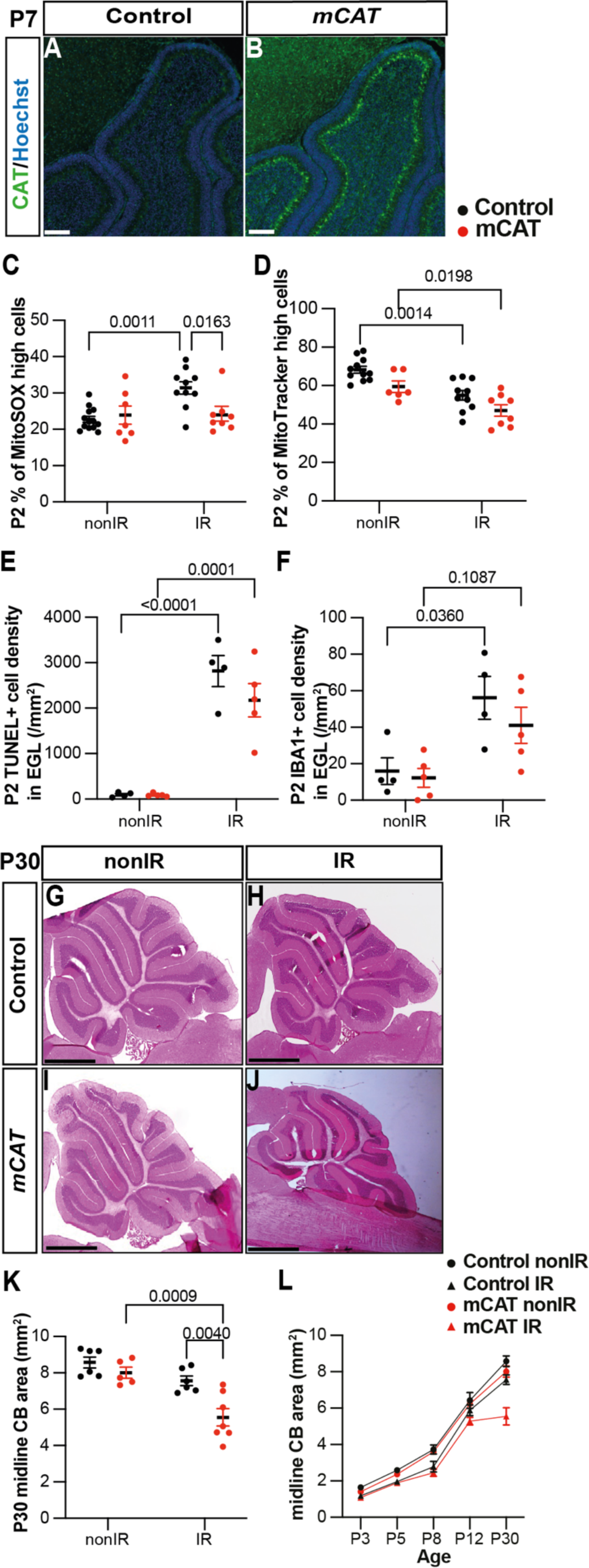
Reduction of ROS impairs adaptive reprogramming and cerebellar repair. **(A, B)** IHC staining of medial sagittal cerebellar sections for human catalase in control (A) AND *mCAT/+* mice (B) at P7. Nuclei were counterstained with Hoechst (blue). Similar staining was seen in four *mCAT/+* mice. **(C)** Quantification of MitoSOX high expression at P2 in control and *mCAT/+* cerebella, with and without irradiation at P1 (Two-way ANOVA, F_(1,34)_=6.768, p=0.0136). **(D)** Quantification of MitoTracker high expression at P2 in control and *mCAT/+* cerebella, with and without irradiation at P1 (Two-way ANOVA, F_(1,31)_=25.06, p<0.0001). **(E)** Quantification of TUNEL+ cell density in the EGL at P2 in control and *mCAT/+* cerebella, with and without irradiation at P1 (Two-way ANOVA, F_(1,14)_=87.56, p<0.0001). **(F)** Quantification of IBA1+ cell density in the EGL at P2 in control and *mCAT/+* cerebella, with and without irradiation at P1 (Two-way ANOVA, F(1,14)=15.58, p=0.0015). **(G-J)** Hematoxylin and eosin staining on mid-sagittal sections of P30 control and *mCAT/+* cerebellum with or without irradiation. **(K)** Quantification of P30 cerebellar mid-sagittal section area in controls and *mCAT/+* nonIR and IR mice (Two-way ANOVA, F_(1,20)_=11.82, p=0.0026). **(L)** Graph showing the average area of mid-sagittal cerebellar sections at P3, P5, P8, P12 and P30 in control and *mCAT/+* non-irradiated and irradiated mice. Detailed statistics are shown in Supplementary Figure 4. EGL, External granular layer; P, postnatal day; nonIR, non-irradiated; IR, irradiated. Scale bar: A and B: 100µm, F-I: 1mm. Significant *Tukey’s post hoc* multiple comparison tests are shown in the figures and data are represented as mean ± SEM.

We next determined the regenerative efficiency of *mCAT/+* mice by analyzing the area of sections of cerebella from nonIR and IR P30 *mCAT/+* mice compared to littermate controls. Strikingly, the cross-sectional area of the medial cerebellum (vermis) of IR *mCAT/+* adult mice was significantly reduced compared to IR controls (p=0.0040, n≥6) (Fig. 4G-K). Analysis of cerebellar area across ages (P3, 5, 8 and 12) revealed that the vermis sectional area of the IR *mCAT/+* cerebella was only significantly reduced at P30 compared to IR controls, however at P12 it was reduced in *mCAT/+* IR cerebella compared to *mCAT/+* nonIR mice, whereas it was not significantly different between control IR and control nonIR mice (Fig. 4L, Supplementary Figure 4 K-N). These results indicate that cerebellar growth begins to be reduced at P12 in *mCAT/+* mice following injury. Thus, a reduction in ROS at the time of cell death in the EGL leads to a diminution of cerebellar recovery.

### Reduced regeneration in *mCAT* mice is associated with reduced adaptive reprogramming at P5

Given that a decrease in ROS following EGL injury reduces regeneration of the neonatal cerebellum, we determined whether specific stages of the adaptive reprogramming process are altered in *mCAT*/+ mice compared to controls. First, we analyzed the replenishment of the EGL by BgL-NEPs in vermis lobules 3-5, since our previous work showed that these lobules have a prominent defect. Interestingly, we found that although the thickness of the EGL in IR mice of both genotypes was similarly reduced compared to nonIR mice at P5 (p=0.0041 control and p=0.0005 *mCAT*/+; n≥4) by P8 the control EGL was a similar thickness to the control nonIR whereas the thickness of the *mCAT*/+ IR EGL was significantly reduced compared to *mCAT*/+ nonIR mice (p=0.035, n=4)(Figure 5A, B). A key regenerative process that contributes to the expansion of the EGL following injury is the migration of BgL-NEPs to the EGL. Strikingly, the density of CFP+ cells in the EGL (mainly BgL-derived NEPs) was significantly decreased in *mCAT*/+ IR mice compared to control IR mice at P5 (p=0.0002, n=4) but not P3 (Figure 5C-G, Supplementary Figure 5A). Furthermore, the density of NEPs (CFP+ or SOX2+ cells) in the BgL was significantly decreased at P5 in *mCAT*/+ IR cerebella compared to controls (p=0.0010, n≥5) but not at other stages (Figure 5H, Supplementary Figure 5B-D). These results indicate that BgL-NEPs have a blunted response to EGL injury, and therefore do not fully expand and contribute to the replenishment of GCPs in the EGL after irradiation.

**Figure 5:**
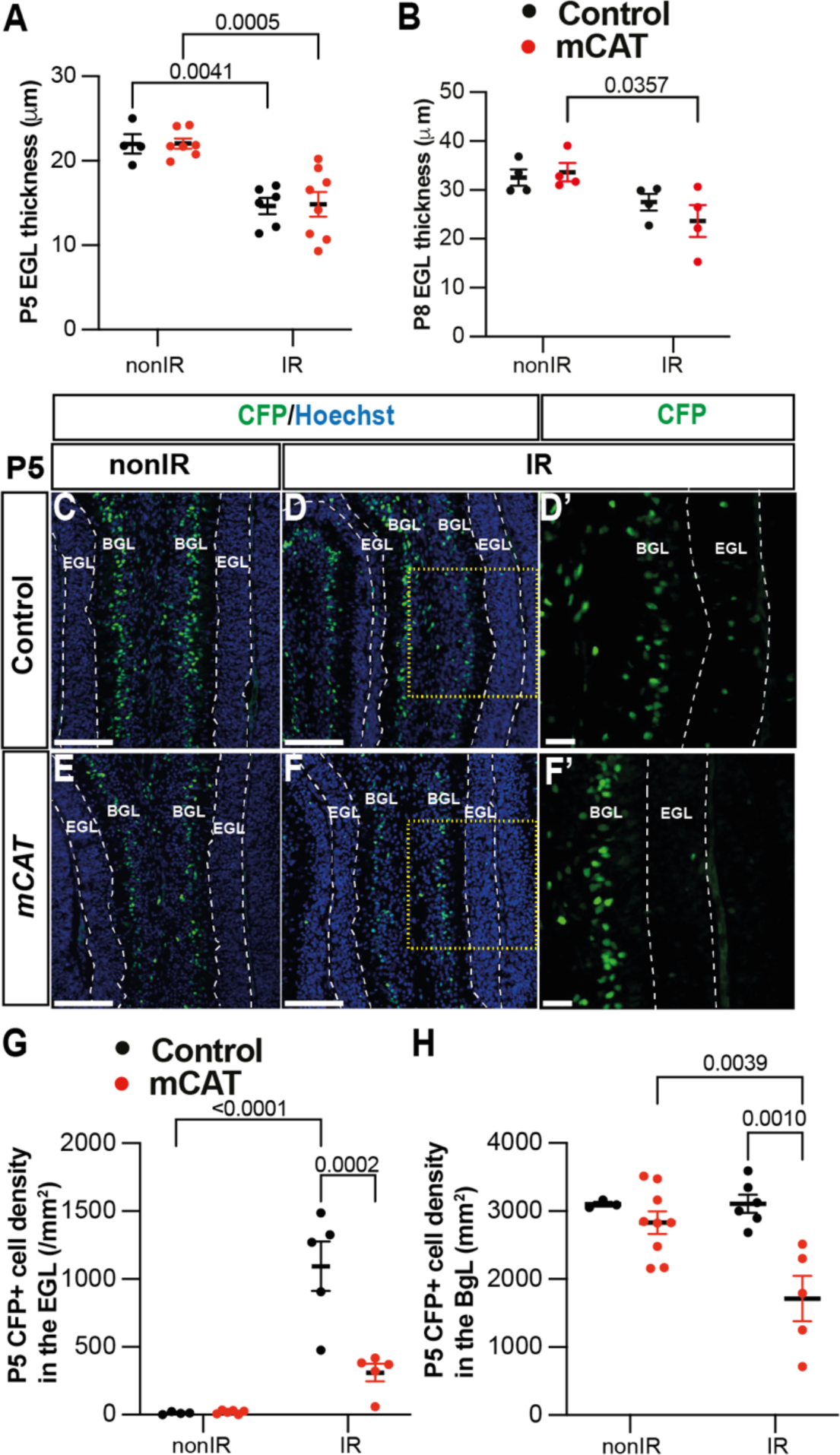
Reduced ROS impairs expansion of BgL-NEPs and their recruitment to the EGL after injury. **(A, B)** Quantification of EGL thickness at P5 (Two-way ANOVA, F_(1,21)_=36.64, p<0.0001) (A) and P8 (Two-way ANOVA, F_(1,12)_=11.34, p=0.0056)(B) in lobules 3-5 of *Nes-Cfp/+* control and *Nes-Cfp/+; mCAT/+* mutant mice with and without irradiation at P1 **(C-F)** IHC staining of medial sagittal cerebellar sections showing expression of CFP (green) in the lobules 4/5 of *Nes-Cfp/+* control and *Nes-Cfp/+; mCAT/+* mutant mice at P5. Nuclei were counterstained with Hoechst (blue). (D’) and (F’) show a high-power images of the yellow boxed area in the single channel CFP. EGL is delineated by the dashed white lines. **(G, H)** Quantification of CFP+ cell density in the EGL (Two-way ANOVA, F_(1,19)_=5.192, p=0.0359) (G) and BgL (Two-way ANOVA, F_(1,17)_=6.191, p=0.0223) (H) at P5 in *Nes-Cfp/+* control or *Nes-Cfp/+; mCAT/+* mutant non-irradiated and irradiated mice. EGL, External granular layer; BgL, Bergmann glia Layer; P, postnatal day; nonIR, non-irradiated; IR, irradiated. Scale bar: D-F: 100µm. Significant *Tukey’s post hoc* multiple comparison tests are shown in the figures and data are represented as mean ± SEM.

### Microglia likely contribute to one aspect of adaptive reprogramming

Given that several steps in adaptive reprogramming were decreased specifically at P5 in *mCAT*/+ IR cerebella, we asked whether microglia/macrophages could be involved in any of the processes. We first determine the density of IBA1+ cells in the EGL and WM of vermis lobules 3-5 at P5 in nonIR and IR mice of both genotypes. As expected, the density of microglia in the EGL was very low in the nonIR control and *mCAT/+* cerebella (Figure 6A-E). Interestingly, whereas control IR mice had a similar number of IBA1+ cells in the WM as nonIR mice of both genotypes at P5, the *mCAT*/+ IR mice had a lower density of microglia/macrophages in the WM compared to control IR mice at P5 (p=0.0012, n≥3) (Figure 6A-D, F). This result raised the question of whether macrophages/microglia play a role in adaptive reprogramming.

**Figure 6:**
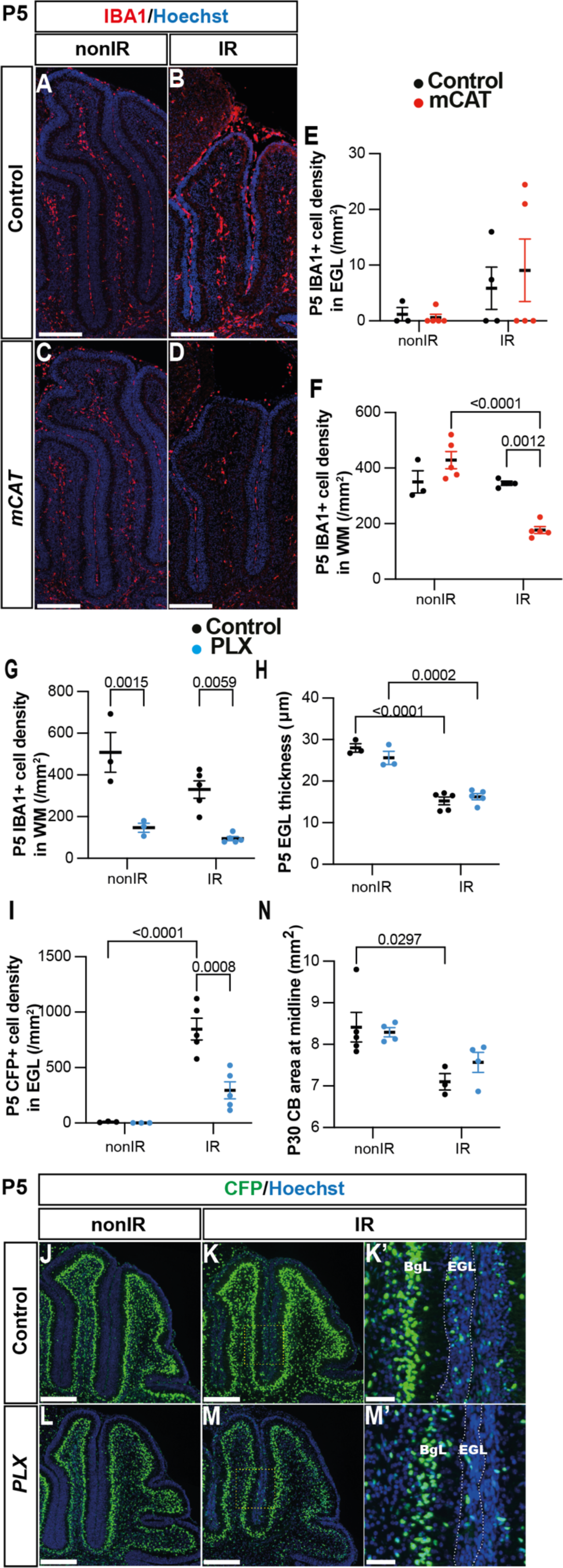
Microglia promote recruitment of NEPs to the EGL during cerebellar adaptive reprogramming after injury. **(A-D)** IHC staining of medial sagittal cerebellar sections for IBA1 (red) in control and *mCAT/+* mice at P5. Nuclei were counterstained with Hoechst (blue). **(E, F)** Quantification of IBA1+ cell density in the external granular layer (E) and white matter (Two-way ANOVA, F_(1,13)_=24.74, p=0.0003) (F) at P5 on midsagittal sections of lobules 3-5 in the cerebellum of control and *mCAT/+* animals, with or without irradiation. **(G-J)** IHC staining of medial sagittal cerebellar sections at P5 for CFP (green) in lobules 4/5 of *Nes-Cfp/+* mice treated with PLX5622 or control DMSO with or without irradiation. Nuclei were counterstained with Hoechst (blue). (H’) and (J’) show a high-power image of area indicated by yellow boxes. EGL is delineated by the white dashed lines. **(K)** Quantification of IBA1+ cell density in the white matter at P5 on mid-sagittal sections in lobules 3-5 of *Nes-Cfp/+* mice treated with PLX5622 or control DMSO, with or without irradiation (Two-way ANOVA, F_(1,12)_=42.40, p<0.001). **(L)** Quantification of EGL thickness at P5 in lobules 3-5 of *Nes-Cfp/+* mice treated with PLX5622 or control DMSO with or without irradiation (Two-way ANOVA, F_(1,12)_=109.5, p<0.001). **(M)** Quantification of CFP+ cells density in the EGL at P5 on mid-sagittal sections in lobules 3-5 of *Nes-Cfp/+* mice treated with PLX5622 or control DMSO with or without irradiation (Two-way ANOVA, F_(1,12)_=10.62, p=0.0068). **(N)** Measurement of cerebellar mid-sagittal section area at P30 in controls or mice treated with PLX, with or without irradiation at P1 (Two-way ANOVA, F_(1,12)_=13.29, p=0.0034). EGL, External granular layer; WM, White matter; P, postnatal day; nonIR, non-irradiated; IR, irradiated. Scale bar: A-D and G-J: 250 µm. Significant *Tukey’s post hoc* multiple comparison tests are shown in the figures and data are represented as mean ± SEM.

We, therefore, tested whether reducing the density of IBA1+ microglia/macrophages after birth would alter adaptive reprogramming at P5 or cerebellar regeneration at later stages. Since macrophages and cerebellar microglia are dependent on colony stimulating factor 1 (CSF1) for their survival, we administered PLX5622, a small molecule inhibitor of CSF receptor 1 (CSFR1), to pups every day from P0-5 (PLX treatment) (Kana et al., 2019, Tan et al., 2021). As expected, IBA1+ cells were significantly decreased in the cerebellum of PLX-treated mice at P5 compared to their controls, both nonIR and IR (p=0.0015 and p=0.0059, n=3 and n=5, respectively) (Figure 6G, Supplementary Figure 6B-E). The thickness of the EGL was not significantly altered at P5 in PLX-treated IR mice compared to IR controls (Figure 6H). Interestingly, similar to *mCAT/+* mice, the density of Nes-CFP+ cells in the EGL was significantly decreased in PLX-treated IR mice compared to IR controls at P5 (p=0.0008, n=5) (Figure 6I-M). In contrast, the density of SOX2+ cells in the BgL, corresponding to the gliogenic BgL-NEPs, were unchanged in the PLX-treated and control mice, whether irradiated or not, suggesting that the decrease in expansion of BgL-NEPs caused by ROS is not mediated by microglia (Supplementary Figure 6F). When mice were treated with PLX from P0-8 and allowed to age to P30, we found the cerebellar vermis section area was not decreased in PLX-treated IR mice compared to IR controls (Figure 6N). Thus, reducing the density of IBA1+ microglia/macrophages in neonatal mice reduces the recruitment of Nes-CFP+ cells to the EGL at P5, but does not have a long-term significant impact on regeneration of the cerebellum.

## Discussion

We demonstrate that a transient increase in ROS signaling after cerebellar injury to the EGL is critical for adaptive reprogramming and full recovery of cerebellar growth. ROS likely acts as an alarm signal shortly after injury. scRNA-seq at P1-5 and bulk ATAC-seq at P2 of NEPs following targeted irradiation at P1 revealed a rapid increase in transcriptional and epigenetic changes associated with upregulation of ROS and stress related pathways in NEPs that peaked at P2. A transient upregulation in ROS at P2 was confirmed using flow cytometry and found to correlate with the timing of cell death in the EGL one day after irradiation. By reducing mitochondrial ROS levels across all cell types at P2 using an *mCAT* transgene, we uncovered that ROS is required for several steps of adaptive reprogramming of BgL-NEPs. In addition, we found that microglia are reduced in injured *mCAT/+* pups, which is consistent with prior evidence that ROS can trigger immune cell recruitment in other systems (Kim et al., 2010, Mehl et al., 2022). Moreover, temporary depletion of microglia caused a reduction in the number of NEPs that migrate into the EGL at P5 following injury at P1, but no long-term reduction of cerebellar size. Thus, we identified key transcriptomic and epigenomic changes in cerebellar NEPs upon GCP ablation at birth and discovered roles for the tissue microenvironment, especially ROS and a more limited role of microglia during neonatal cerebellum regeneration.

scRNA-seq analysis of NEPs from nonIR (P1-3, P5) and nonIR (P2, P3, P5) showed an increase in genes associated with stress responses after irradiation in both the gliogenic and neurogenic subpopulations, but not in the GCPs. Furthermore, an increase in ROS signaling was detected one day after injury (P2) in all three lineages. The injury-induced ROS and stress-related gene signatures were not observed in the later P3+5 gliogenic- and neurogenic-NEPs, suggesting that the increase in ROS levels is an early injury induced signal affecting the NEP transcriptome. Our bulk ATAC-seq data generated at P2 revealed that the open chromatin regions in the IR NEPs were enriched for transcription factor binding motifs related to ROS signaling and stress induced transcription factors such as FOXO3 and AP1 (Table S6). Thus, cerebellar injury during development induces transcriptional and epigenomic signatures in the NEPs and ROS signaling could be a key driver of NEP adaptive reprogramming. Our results are in line with other regeneration systems where a temporary increase in ROS is observed upon cell death or injury and is considered to be a DAMP (Niethammer, 2016).

Interestingly, although both gliogenic- and neurogenic-NEPs showed induction of cellular stress related genes upon injury, upregulation of ROS signaling and related genes appeared greater in the *Hopx*-expressing gliogenic NEPs that undergo adaptive reprogramming. Whether the upregulation of ROS signaling in the BgL-NEPs is due to BgL-NEPs being in proximity to the dying GCPs after injury or their direct contact due to their radial projections remains to be determined. The ability of BgL-NEPs to respond to GCP death via upregulating ROS signaling and impaired regeneration upon reduction of ROS levels, shows that ROS signaling is involved in triggering adaptive reprogramming upon injury.

The cellular composition of the neonatal cerebellum is dramatically different from the adult. During the early postnatal period, we found that astrocytes in the WM below the lobules are the first to initiate GFAP expression at P2 and that by P5 all astrocytes express a high level of GFAP. In contrast, Bg express a low level of GFAP at P5 and reach a high level by P8. Interestingly, we found that injury leads to an increase in the level of GFAP expression in each type of astroglia when they first initiate expression, deep WM astrocytes at P2 and Bg (and all astrocytes) at P5. Once adaptive reprogramming is nearing completion (P8), the GFAP levels remained similar in all astroglia in nonIR and IR cerebella. These results suggest that the neonatal astrocytes respond to injury differently than in the adult where GFAP upregulation is observed immediately after injury (Burda and Sofroniew, 2014), since Bg and astrocytes in the lobules have a delayed response to injury with GFAP not being upregulated for several days. Furthermore, at birth the microglia have not fully expanded in number in the cerebellum and previous scRNA-seq showed that the neonatal and adult microglia are transcriptionally distinct (Hammond et al., 2019). Perhaps neonatal microglia are anti-inflammatory and pro-regenerative upon injury in neonates, in contrast to adult microglia where upon traumatic brain injury they can inhibit regeneration (Donat et al., 2017). Collectively, the differences in glial responses to injury in the neonatal cerebellum and adult brain likely contributes to the permissiveness of the neonatal cerebellum to regeneration. Details of the molecular changes that the neonatal cerebellar microglia/macrophages and astrocytes undergo upon GCP injury and their molecular crosstalk with NEPs remain to be determined.

We demonstrated the significance of ROS activation in the NEP reprogramming process using an *mCAT* transgenic mouse line in which human Catalase (CAT) is expressed in mitochondria, a protein that can lead to a reduction in hydrogen peroxide (H_2_O_2_) and thus lower ROS. However, in *mCAT* neonatal cerebella we found that the percentage of MitoSOX high cells are comparable to control mice under nonIR conditions. Importantly, however, one day after irradiation at P1 when the percentage of MitoSOX high cells increases in control IR mice, the *mCAT* transgene reduces the percentage of MitoSOX high cells, such that it remains at the baseline nonIR *mCAT* level.

Therefore, human CAT expression in mitochondria in this model inhibits the injury-induced increase in ROS levels without affecting the homeostatic production of superoxide. Of possible relevance, in this mouse model the observed change in ROS levels is likely global, impacting all cell types. The specific impact of increased mCAT in BgL-NEPs or microglia on their recruitment and function after an injury remains to be determined with new cell type-specific tools.

Our experiments depleting microglia using PLX5622 indicate that microglia/macrophages are involved in the regeneration of the EGL following irradiation. Previous studies have demonstrated a dual role of microglia in promoting and inhibiting regenerative processes within the nervous system (Lee et al., 2021, Wang et al., 2020). Our data support the idea that microglia are involved in the adaptive reprogramming of NEPs to GCPs by promoting their replenishment of GCPs in the EGL. While the direct mechanisms remain to be discovered, given the small size of the lobules and disruption of the cytoarchitecture after injury it might be possible for secreted factors from the white matter microglia to reach the BgL NEPs. Alternatively, there could be a relay system through an intermediate cell type closer to the microglia. PLX-treatment for 8 days after irradiation did not reduce the later growth of the injured cerebellum. One possibility is that after the cessation of PLX administration, regeneration proceeds normally. Additionally, regenerative processes that act in parallel to microglial signaling are likely required.

Collectively, we have delineated the spatiotemporal cellular changes in the cerebellar glial microenvironment upon ablation of GCPs at birth and highlight ROS signaling as a key stimulator of adaptive reprogramming of NEPs. The details of how DAMP-glia-progenitor crosstalk is orchestrated remains to be untangled. Understanding how microenvironmental responses shape repair processes is a crucial first step towards developing strategies to promote regeneration.

## Acknowledgements

We thank past and present members of the Joyner laboratory for discussions and technical help. We would like to thank Dr Ronan Chaligne and his team for their support in the multiplexed scRNA-seq experiments. We are grateful to the MSKCC Animal Imaging Core, Flow cytometry Core, Center for Comprehensive Medicine and Pathology, Integrated Genomics Operation, Single-cell Analytics and Innovation Laboratory and Epigenetics Computational Laboratory teams for technical services and support. An XRad 225Cx Microirradiator was purchased by support from a Shared Resources Grant from the MSKCC Geoffrey Beene Cancer Research Center. We gratefully acknowledge the support of the Gurdon Institute Scientific Computing Facility.

This work was supported by grants from the NIH to ALJ (R01NS092096) and NSB (NINDS K99 NS112605-01). Additional funding was provided to ALJ from an NCI Cancer Center Support Grant (CCSG, P30 CA08748) and the Cycle for Survival, to SEN from a Francois Wallace Monahan Fellowship; to NSB from a Wellcome Career Development Award (227294/Z/23/Z), Royal Society grant (RGS\R1\231143) and Cambridge Stem Cell Institute Seed Funding, and to JBC from a University of Cambridge School of Biological Sciences DTP PhD Studentship and Peter and Emma Thomsen’s Scholarship (1051). Gurdon Institute is supported by a Wellcome Core Grant (203144) and CRUK Grant (C6946/A24843).

---

**Supplementary Figure 1.**
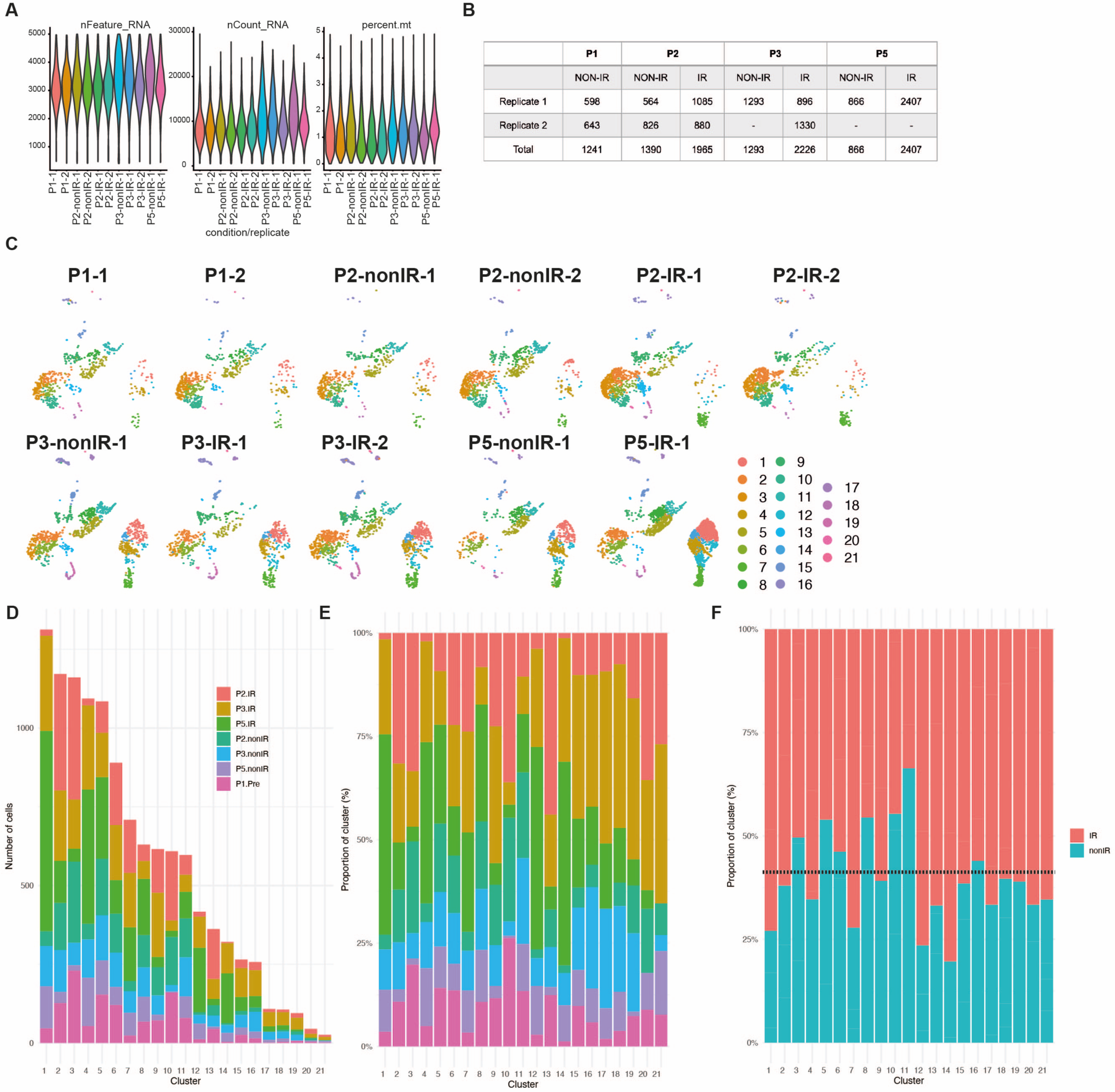
scRNA-seq quality metrics and number of cells sequenced in each condition and biological replicate. **(A)** Violin plots showing the number of features, RNA and percent mitochondrial RNA count across the biological replicates of the scRNA-seq data set after filtering the bad quality cells (cells were filtered out where number of detected genes was ≤ 1500, the number of detected transcripts was ≥ 40,000 and mitochondrial gene percentage ≥ 5%). **(B)** Number of cells from each replicate and condition used for downstream analyses after filtering. **(C)** UMAPs showing the distribution of cells across different clusters based on the samples. **(D-E)** Number and proportion of cells from different ages and conditions in each cluster. **(F)** Proportion of total nonIR (including P1) and IR cells in each cluster. Dotted line represents the expected ratio between the nonIR and IR cells.

**Supplementary Figure 2:**
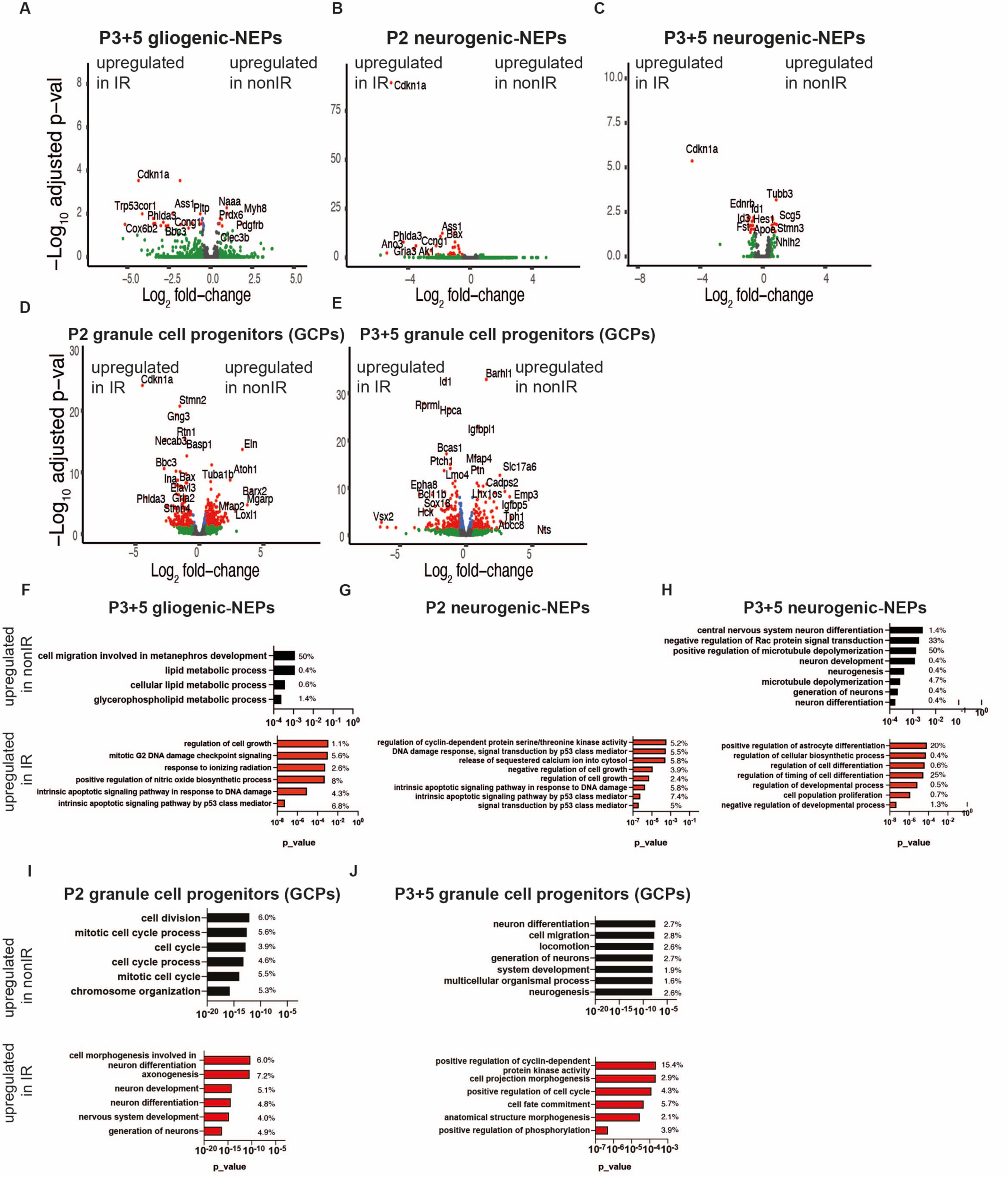
Injury induces distinct transcriptional changes in NEP subtypes and GCPs during adaptive reprograming. **(A-E)** Volcano plot showing differentially expressed genes in the P3+5 gliogenic-NEPs (A), P2 and P3+5 neurogenic NEPs (B, C) and P2 or P3+5 GCPs (D, E) (red: adjusted p-value≤0.05, log_2_fold-change=1, Table S2). **(F-J)** Some of the significant GO terms associated with differentially expressed genes in the P3+5 gliogenic-NEPs (F), P2 or P3+5 neurogenic NEPs (G, H) and P2 or P3+5 GCPs (I, J) that were either upregulated in nonIR (top panel) or IR (bottom panel) (p-value≤0.05).

**Supplementary Figure 3:**
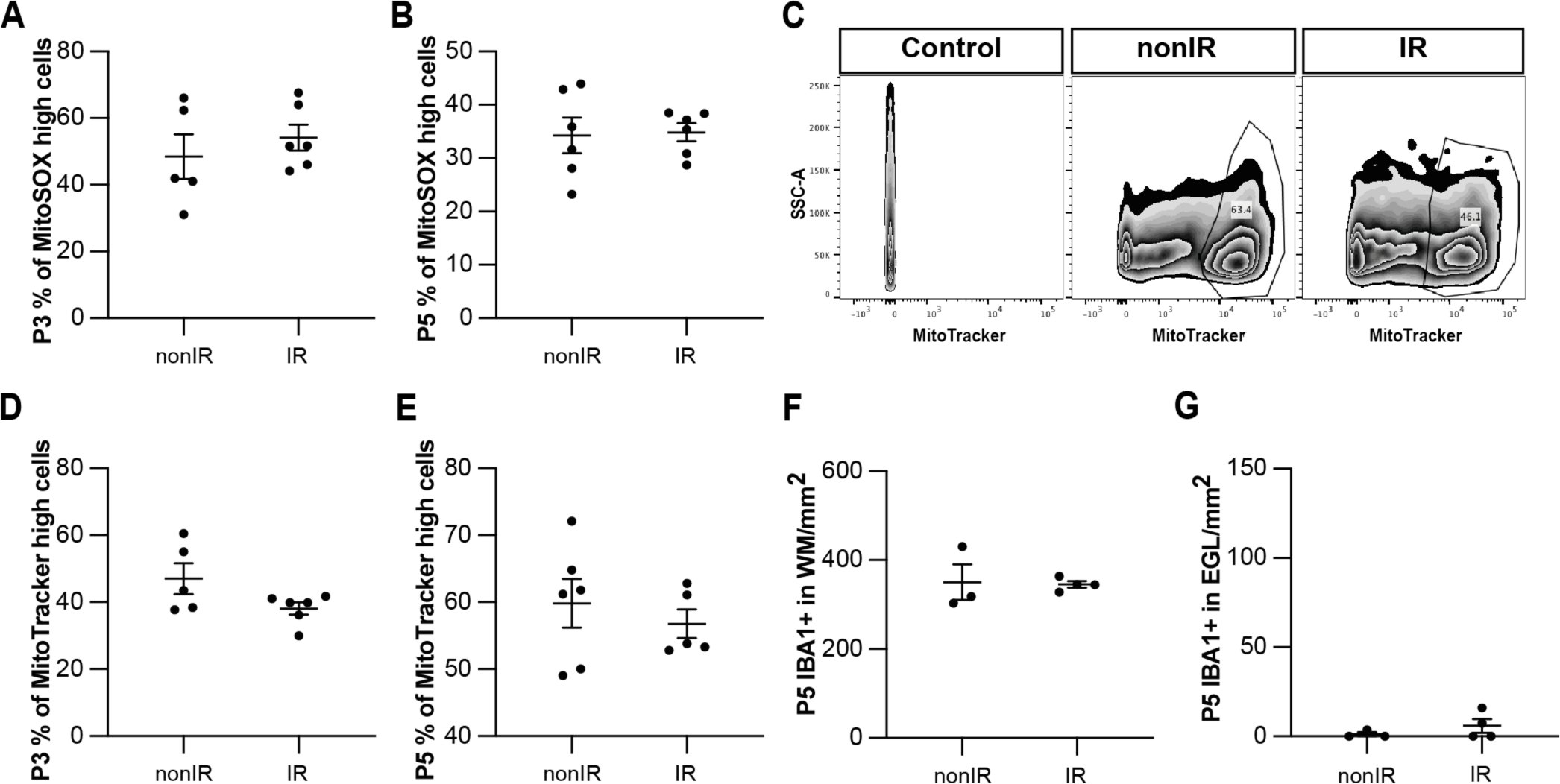
Irradiation of cerebella at P1 results in increased superoxide production and cell death and recruitment of microglia to the EGL that peaks at 24h. **(A, B)** Quantification of high MitoSOX expression in nonIR and IR cerebella at P3 (A) and P5 (B). **(C)** Examples of flow cytometry analysis of mitochondria at P2 from nonIR and IR cerebella using MitoTracker dye. Gating determined the top 90% MitoTracker signal (MitoTracker high cells). **(D, E)** Quantification of MitoTracker high expression in nonIR and IR cerebella at P3 (D) and P5 (E). **(F)** Quantification of IBA1+ cell density in the WM at P5 in lobules 3-5 of nonIR and IR mice. **(G)** Quantification of IBA1+ cell density in the EGL at P5 in lobules 3-5 of nonIR and IR mice. EGL, External granular layer; WM, White matter; P, postnatal day; nonIR, non-irradiated; IR, irradiated. All statistical significance was determined using an unpaired t-test and data are represented as mean ± SEM.

**Supplementary Figure 4:**
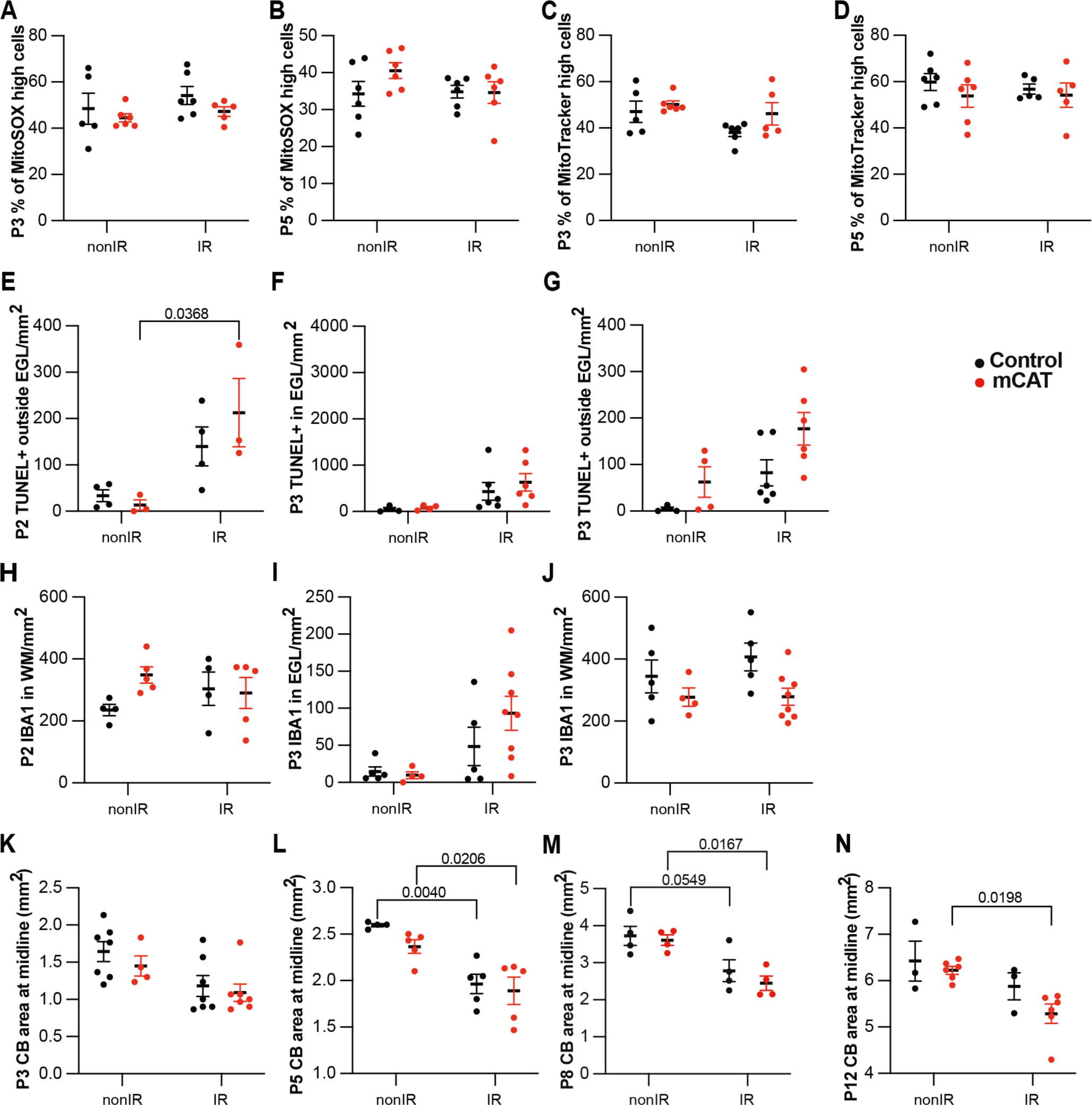
Reduction of ROS impairs adaptive reprogramming and cerebellar repair. **(A, B)** Quantification of high MitoSOX expression at P3 (A) and P5 (B) in control and *mCAT/+* cerebella, with and without irradiation at P1. **(C, D)** Quantification of MitoTracker high expression at P3 (C) and P5 (D) in control and *mCAT/+* cerebella, with and without irradiation at P1. **(E-G)** Quantification of TUNEL+ cell density outside EGL at P2 (Two-way ANOVA, F_(1,10)_=14.20, p=0.0037) (E), at P3 (G), and in the EGL at P3 (F) in lobules 3-5 of nonIR and IR mice. **(H-J)** Quantification of IBA1+ cell density in WM at P2 (H), at P3 (J), and in the EGL at P3 (I) in lobules 3-5 of nonIR and IR mice. (**K-N)** Quantification of cerebellar midsagittal section area in controls and *mCAT/+* nonIR and IR mice at P3 (K), P5 (Two-way ANOVA, F_(1,15)_=28.52, p<0.001) (L), P8 (Two-way ANOVA, F_(1,12)_=21.21, p=0.0006) (M) and P12 (Two-way ANOVA, F_(1,14)_=9.682, p=0.0077) (N). EGL, External granular layer; WM, White matter; P, postnatal day; nonIR, non-irradiated; IR, irradiated. Significant *Tukey’s post hoc* multiple comparison tests are shown in the figures and data are represented as mean ± SEM.

**Supplementary Figure 5:**
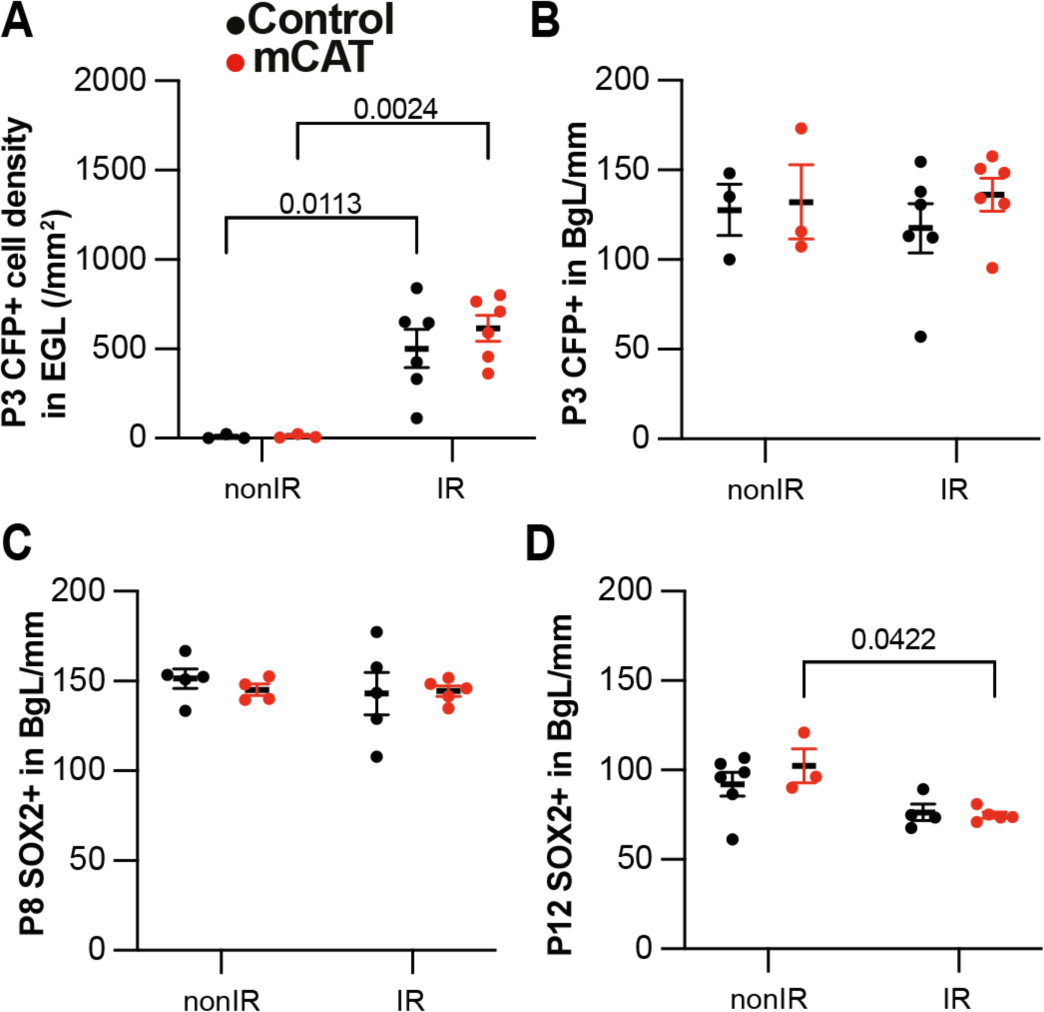
Reduced ROS impairs expansion of BgL-NEPs and their migration to the EGL after injury. **(A)** Quantification of CFP+ cell density in the EGL at P3 in *Nes-Cfp/+* control or *Nes-Cfp/+ mCAT* mutant nonIR and IR mice. (Two-way ANOVA, F_(1,14)_=33.77, p<0.0001). **(B)** Quantification of CFP+ cell normalized on BgL length at P5 in *Nes-Cfp/+* control or *Nes-Cfp/+ mCAT* mutant nonIR and IR mice. **(C, D)** Quantification of SOX2+ NEP cell density on BgL length at P8 (C) and P12 (Two-way ANOVA, F_(1,12)_=12.50, p=0.0033) (D) in control or *mCAT* mutant nonIR and IR mice. EGL, External granular layer; BgL, Bergmann glia Layer; P, postnatal day; nonIR, non-irradiated; IR, irradiated. Significant *Tukey’s post hoc* multiple comparison tests are shown in the figures and data are represented as mean ± SEM.

**Supplementary Figure 6:**
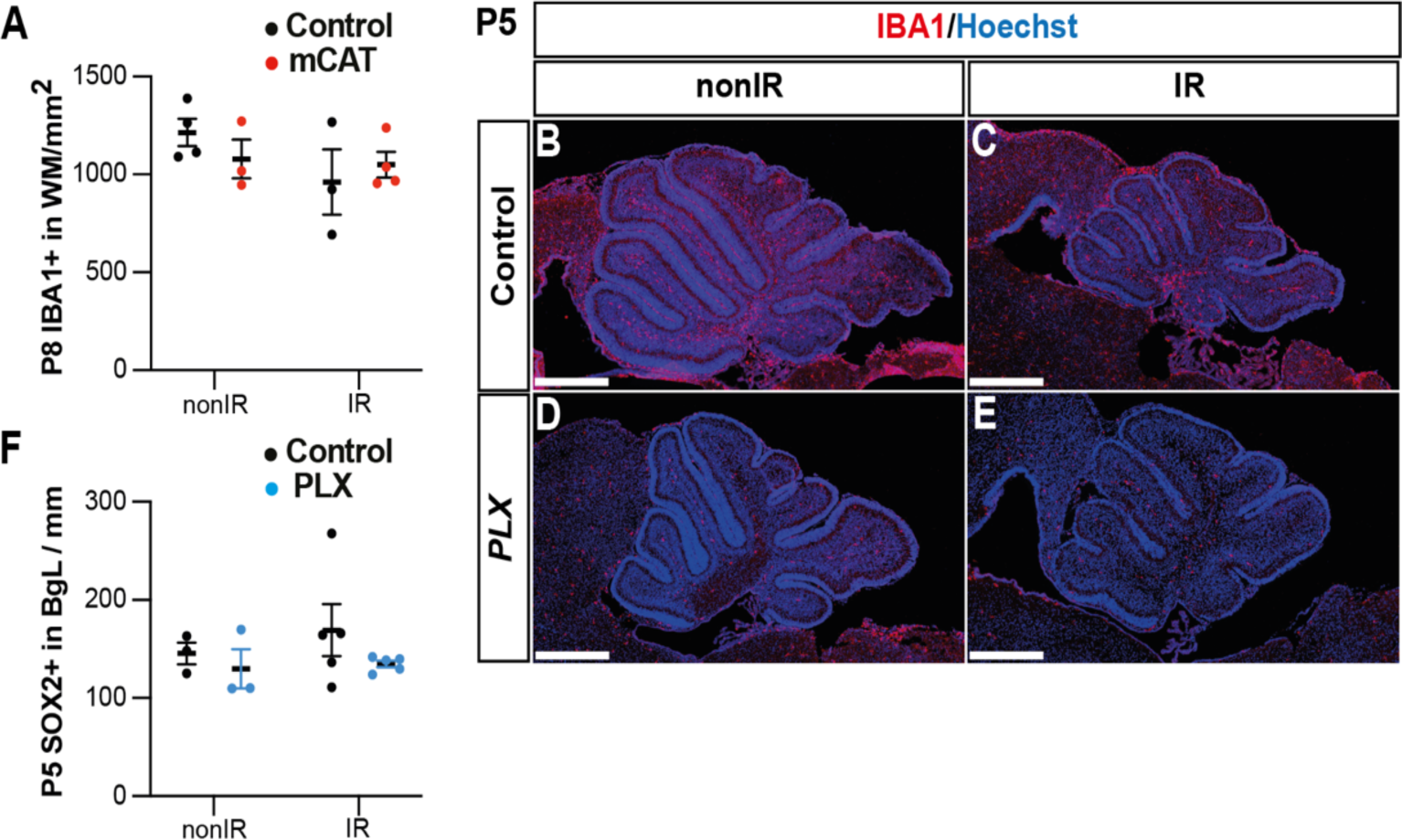
Microglia promote recruitment of NEPs to the EGL during cerebellar adaptive reprogramming after injury. **(A)** Quantification of IBA1+ cell density in the WM at P8 in control or *mCAT* mutant nonIR and IR mice. **(B-E)** Immunostaining of medial sagittal cerebellar sections at P5 showing expression of IBA1 (red) in mice treated with PLX5622 or control DMSO, with or without irradiation. (F) Quantification of SOX2+ NEP cell density in the BgL at P5 in control or PLX treated nonIR and IR mice. BgL, Bergmann glia Layer; WM, White Matter; P, postnatal day; nonIR, non-irradiated; IR, irradiated. Scale bar: 500 µm. Data are represented as mean ± SEM.

## Supplementary Tables

**Table S1:** List of antibodies and related information.

| Antigen | Species | Concentration | References | Source |
| --- | --- | --- | --- | --- |
| Catalase | Rabbit | 1 to 100 | 01-05-030000 | Athens Research & Technology |
| GFAP | Chicken | 1 to 500 | ab4674 | Abcam |
| GFP | Rat | 1 to 1000 | 440484 | Nacalai Tesque |
| IBA1 | Rabbit | 1 to 500 | 019-19741 | Wako Chemicals |
| SOX2 | Goat | 1 to 500 | AF2018 | R&D System |

**Table S2.** Marker genes expressed by cluster in scRNA-seq dataset (irradiated at P1 (IR; P2, P3, P5) or non-irradiated (nonIR; P1, P2, P3, P5). pct1: % cells in a cluster that express the gene; pct2: % cells that express the gene outside the given cluster.

**Table S3.** Pseudobulk differential expression analysis between nonIR and IR gliogenic-NEPs (*Hopx+,* clusters 2, 3, 6, 10), neurogenic-NEPs (*Ascl1+,* clusters 5, 8, 11) and GCPs (*Atoh1+,* clusters 1,4,7,12,14) at P2, or at P3 and P5 (P3+5).

**Table S4.** GO Term analyses of differentially expressed genes (Table S3) of nonIR and IR gliogenic-NEPs (*Hopx+,* clusters 2, 3, 6, 10), neurogenic-NEPs (*Ascl1+,* clusters 5, 8, 11) and GCPs (*Atoh1+,* clusters 1,4,7,12,14) at P2, or at P3 and P5 (P3+5).

**Table S5.** Differentially open peaks at P2 identified by bulk ATAC-seq from nonIR and IR NEPs.

**Table S6.** Motif analysis of regions with increased accessibility in IR NEPs compared to the nonIR at P2.

